# VikNGS: A C++ Variant Integration Kit for Next Generation Sequencing association analysis

**DOI:** 10.1101/504381

**Authors:** Zeynep Baskurt, Scott Mastromatteo, Jiafen Gong, Richard F. Wintle, Stephen W. Scherer, Lisa J. Strug

**Affiliations:** Program in Genetics and Genome Biology, Research Institute, The Hospital for Sick Children, Toronto, Ontario, M5G0A4, Canada; The Centre for Applied Genomics, The Hospital for Sick Children, Toronto, Ontario, M5G0A4, Canada; Division of Biostatistics, Dalla Lana School of Public Health, University of Toronto, Ontario, M5T3M7, Canada; McLaughlin Centre and Department of Molecular Genetics, University of Toronto, Toronto, ON, M5G 0A4, Canada.

## Abstract

**Motivation:** Integration of next generation sequencing data (NGS) across different research studies can improve the power of genetic association testing by increasing sample size and can obviate the need for sequencing controls. Unfortunately, if differential genotype uncertainty across studies is not accounted for, combining data sets can also produce spurious association results. The robust variance score statistic (RVS) for genetic association of rare and common variants has been shown to effectively adjust for bias caused by the differences in read depth in case-control genetic association studies when the two groups were sequenced using different experimental designs. To enable consortium research, the aggregation of several data sets for genetic association analysis of quantitative and binary traits with covariate adjustment is required, and we developed the Variant Integration Kit for NGS (VikNGS) that expands the functionality of RVS (vRVS) for this purpose.

**Results:** VikNGS is a fast and computationally efficient cross-platform software package that provides an implementation for vRVS, as well as conventional rare and common variant genotype-based association analysis approaches. The package includes a graphical user interface that contains power simulation functionality and data visualization tools.

**Availability and Implementation:** The VikNGS package can be downloaded at http://www.tcag.ca/tools/index.html

Documentation can be found at https://VikNGSdocs.readthedocs.io/en/latest/

**Supplementary information:** Supplementary data are available at *Bioinformatics* online.

## 1 Introduction

Genetic association studies have contributed greatly to our understanding of complex traits. However, to achieve the sample sizes necessary to carry out genome-wide association investigations, the last two decades has seen a shift in the design of studies from individual investigator-oriented to collaboration-based research led by large consortia (Austin, Hair, & Fullerton, 2012). This has historically been achieved by combining genome-wide genotype array data across study groups for meta or mega genome-wide association analysis (Tang & Lin, 2014) of common variants. Structural or rare variants also contribute to disease variation, but are less well captured by genotyping arrays (Eichler et al., 2010). Association studies with whole genome sequencing (WGS) enables analysis of the full allele frequency spectrum and the decreasing cost of this technology continues to make it more accessible. Yet collaboration across study groups along with the integration of other publicly available WGS data are required to realize the statistical power necessary for the identification of associated loci.

Depending on resources and the scientific context, different groups may choose different experimental designs for their WGS studies. For example, the UK10K Consortium (TheUK10KConsortium, 2015) aimed to implement WGS on preexisting cohorts where several phenotypes were available. Since sequencing costs scale with read depth, projects with a large number of participants may choose to sequence at a lower read depth (average 6.5x for UK10K). The large sample size and the broad phenotypic information available in the UK10K Consortium data suggests potential as a convenience control group in case-control association studies. The 1000 Genomes Project is another well known example of publicly available low read depth (4x) WGS data that could be used as a control group (The 1000 Genomes Project Consortium, 2015). For sequencing of small patient populations, high read depth designs are more frequently implemented.

From next generation sequencing (NGS) data, confidence in a variant call is dependent on, among other factors, the sequencing technology, read depth, error rate, base calling algorithm, alignment, SNP detection and genotype calling algorithms (Nielsen, Paul, Albrechtsen, & Song, 2011). (Skotte, Korneliussen, & Albrechtsen, 2012) developed a score statistic which accounts for the uncertainty in genotype calls in a given study. Their method provides a score test where genotype calls are substituted by their expected values, and results indicate better control of Type I error and higher power when compared to association methods based on genotype calls. Building on the approach by (Skotte, Korneliussen, & Albrechtsen, 2012), (Yan et al., 2015) proposed to take genotype calling uncertainty into account through a combined score and likelihood ratio test that leverages the advantages of both tests. In their score test they implement an alternative information matrix to that used in (Skotte et al., 2012) which improves power. (Yan et al., 2015) did not investigate the performance of their method when combining data from cohorts sequenced independently, potentially using different sequencing designs, and their method was applied for single variant analysis.

Although not widely appreciated, naively pooling genotype calls from different WGS studies and performing an association test can result in spurious association findings due to the bias introduced by the differences in genotype call uncertainty across experimental designs. Previously, we showed the impact of differential read depth on case-control genetic association studies when the cases and controls were sequenced with high and low read depth, respectively (Derkach, Chiang, et al., 2014). Building on the method proposed in (Skotte et al., 2012) to account for the uncertainty of the genotype calls, we developed a robust variance score statistic (RVS) to adjust the bias resulting from read-depth differences (Derkach, Chiang, et al., 2014). The approach achieves Type I error rate control when publicly available low-read-depth controls are used in case-control association studies with high read-depth sequencing of cases, which has already contributed to novel gene discovery (Luo et al., 2017). When combining WGS from different study groups spurious association findings can also result from failure to adjust for confounders, from unbalanced case-control designs and from the combination of sequence across greater than two groups for association studies. Currently, methodology that addresses these additional considerations does not exist.

(Hu, Liao, Johnson, Allen, & Satten, 2016) also considered the scenario for which RVS was designed, where cases and controls are sequenced at different read depth using different experimental designs. They developed a screening algorithm to estimate variant loci using the read data instead of relying on existing genotype calling approaches. Their score statistic for association has the same form as the RVS (Derkach, Chiang, et al., 2014), but their genotype likelihoods are calculated from their own algorithm instead of obtaining them from the output of a standard genotype calling package. More recently, (Lee, Kim, & Fuchsberger, 2017) proposed a bias correction on the estimate of the log odds ratio when external controls are added to the study and derived a score-type statistic using this bias corrected log odds estimate for single or region based rare variant association tests. This approach only requires allele counts from external groups which is an appealing property but requires an internally sequenced case and control group and assumes only one external control group is to be added.

Here we introduce the Variant Integration Kit for next generation sequencing (NGS), VikNGS, which is a fast and computationally efficient package developed in C++ that enables genetic association testing using NGS data. Given a set of variant calls from an arbitrary number of studies, VikNGS offers a suite of tests that can be used to identify associated variants. VikNGS builds on the RVS framework (Derkach, Chiang, et al., 2014) and includes an extended version of the methodology (vRVS) which enables integration of sequence across any arbitrary number of data sets that may have been sequenced using different experimental designs and enables common and rare variant association testing for binary or quantitative traits with covariate adjustment. The software also provides conventional common and rare variant association tests with genotype calls, e.g. CAST (Morgenthaler & Thilly, 2007) and SKAT (Wu et al., 2011), as well as power and sample size estimation for study planning. The general workflow for VikNGS is shown in Figure 1. In Section 2, we present the vRVS framework and explain the usage of VikNGS. We demonstrate the robustness of vRVS through a comprehensive simulation study using VikNGS and compare Type I error and power of the vRVS to conventional methods using genotype calls with comparison to the true genotypes used as the gold standard. In Section 3, we apply VikNGS to genetic association studies in Cystic Fibrosis (CF) and compare results to conventional approaches. First, in a proof-of-concept study we integrate NGS from chromosome 7, which contains the CF-causing *cystic fibrosis transmembrane conductance regulator* (*CFTR*; chr7:117,110,017-117,318,718; hg19) (Kerem et al., 1989), for 101 individuals of European descent with CF sequenced at an average read depth of 30x, 1,927 non-CF individuals from the UK10K sequenced at an average read depth of 6.5x and 379 individuals of European descent from the 1000 Genomes Project Phase 1 sequenced at 4x read depth. We then implement VikNGS in a quantitative trait analysis using 1,927 participants from the UK10K at a previously identified CF modifier locus on chromosome 1. This latter analysis demonstrates VikNGS functionality to implement conventional tests and to assess the association evidence of a CF modifier gene with lung function in a non-CF population. In Section 4, we discuss the functionality, strengths and limitations of the vRVS framework and VikNGS, along with aspects for future development.

**Figure 1.**
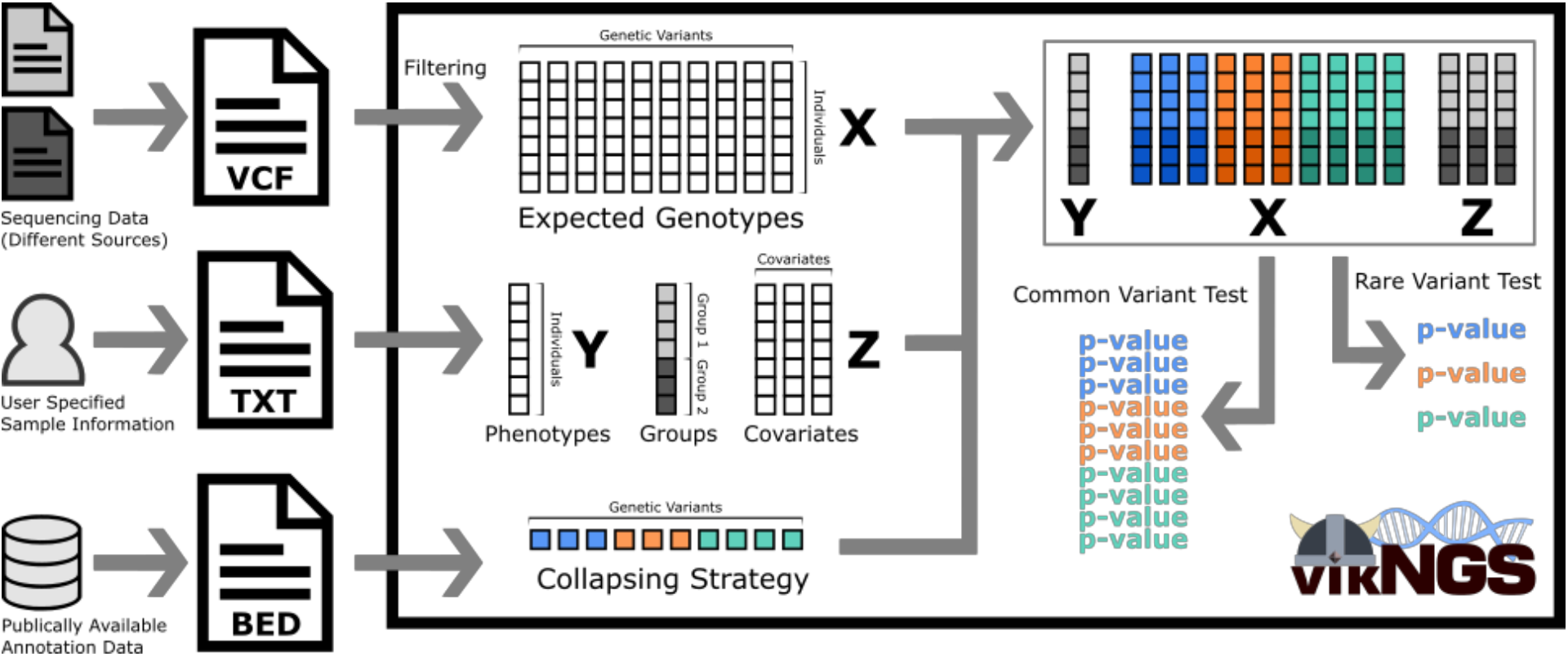
VikNGS workflow for NGS association analysis when sequenced samples from different studies are combined. Expected genotypes are computed from a VCF file. Phenotype, group and covariate information is provided in a tab-separated text file. Optionally, a BED file specifying genomic intervals can be used to define a variant collapsing strategy. VikNGS will parse this data and perform a series of association tests.

## 2 Methods

### 2.1 The vRVS and the score test

Consider the joint likelihood of the observed phenotype, *Y_i_*, and the observed sequencing data, *D_ij_*, for individual *i* at locus *j*, for *i* = 1, …, *n* independent samples,

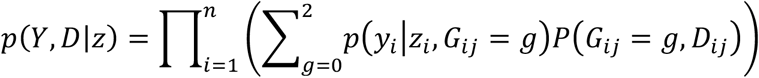

where *Y_i_* depends on *D*_*ij*_ through the unobserved genotype *G*_*ij*_ (taking values 0,1 or 2) and z can be any observed additional covariates. We derive a score statistic based on this model following the approach proposed in (Skotte et al., 2012) and in (Derkach, Chiang, et al., 2014). For case-control studies it is common to use logistic regression and in this case *p*(*y*_*i*_|*z*_*i*_, *G*_*ij*_) = *exp*(*β*_0_ + *β*_1_*g*_*i*_ + *αz*_*i*_)/(1 + *exp*(*β*_0_ + *β*_1_*g*_*i*_ + *αz*_*i*_)). If the phenotype is a normally distributed quantitative trait, then we assume *Y_i_* is distributed as *N*(*β*_0_ + *β*_1_*g*_*i*_ + *αz*_*i*_, *σ*^2^). The score function under *H*_0_: *β*_1_ = 0 is 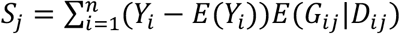 and the corresponding score test statistic is 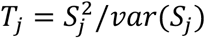 and approximately *χ*^2^ distributed with 1 degree of freedom (Skotte et al., 2012). Note that the estimate of *E*(*Y*_*i*_) is simply 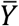 when there are no covariates in the model and *α* can be a vector if the number of covariates is greater than one. *E*(*G*_*ij*_|*D*_*ij*_) is calculated using the expectation formula, 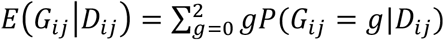 where *P*(*G*_*ij*_|*D*_*ij*_) = *P*(*D*_*ij*_|*G*_*ij*_ = *g*)*P*(*G*_*ij*_ = *g*)/*P*(*D*_*ij*_). We can obtain the conditional probabilities from the output of standard genotype calling packages, such as the variant calling format (VCF) files. The genotype probabilities *P*(*G*_*ij*_ = *g*) can be calculated using an EM algorithm that incorporates the full sample (McKenna et al., 2010).

To briefly summarize how the RVS (Derkach, Chiang, et al., 2014) builds on this model, consider the calculation of the variance of the score function, *var*(*S*_*j*_), which needs to evaluate *var*(*E*(*G*_*ij*_|*D*_*ij*_)). By the law of total variance, we see that *var*(*G*_*ij*_) can be decomposed as *var*(*G*_*ij*_) = *var*(*E*(*G*_*ij*_|*D*_*ij*_) + *E*(*var*(*G*_*ij*_|*D*_*ij*_). When read depth is high, the probability of *G* given *D*, *P*(*G*_*ij*_|*D*_*ij*_), converges to 1. Thus, *E*(*G*_*ij*_|*D*_*ij*_) converges to the true value of genotype *G_ij_* and thus *var*(*E*(*G*_*ij*_|*D*_*ij*_)) converges to the *var*(*G*_*ij*_). However, for low coverage data, *E*(*var*(*G*_*ij*_|*D*_*ij*_))>0 and *var*(*E*(*G*_*ij*_|*D*_*ij*_)) is smaller than *var*(*G*_*ij*_). That is, *var*(*E*(*G*_*ij*_|*D*_*ij*_) is read depth dependent and should be accounted for in the estimation of the variance in the score test. In (Derkach, Chiang, et al., 2014) the scenario where cases sequenced at high read depth were combined with controls sequenced at low read depth was considered, and the variance estimate was shown to be biased with the bias depending on the sample size in each group and the variance difference between the two groups. To overcome this bias, the variance of cases and controls can be estimated separately and when the number of cases is small, one can replace 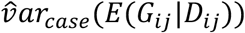 with 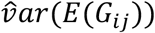 where 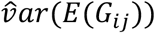 is calculated through the estimated *P*(*G*_*ij*_ = *g*) using the full data set. The variance of the control group is calculated using the sample variance of the low read depth (LRD) group, e.g. 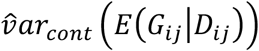. (Derkach, Chiang, et al., 2014) showed that the variance of the smaller group (e.g. case at high read depth (HRD)) has a larger weight on the variance of *S_j_* i.e. 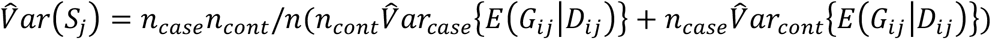, since the sample size for cases (*n_case_*) is smaller than the control group (*n_cont_*), and that this variance estimation procedure is robust, in that it controls Type I error inflation due to differences in read depth between cases and controls.

#### Extensions for Binary Trait Analysis

For combining more than two datasets, if the number of cases is smaller than the number of controls, the variance of the case groups will have a larger effect on the variance of *S_j_*. Suppose there are *K* groups with sample sizes *n*_*k*_, *k* = 1, …, *K*. Using the variance formula 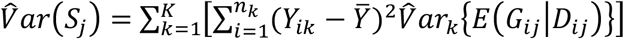 to calculate the variance of the score, *S_j_*, the 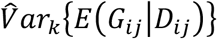 is estimated by 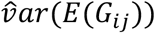 if group *k* is sequenced at HRD and is estimated by the sample variance in group *k* if group *k* is sequenced at LRD. The effectiveness of the RVS is most apparent when the sample size of the HRD groups is smaller than that of the LRD groups. When covariates are added to the model, 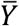 is replaced with 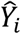, where 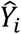 is the fitted values obtained from the regression model under *H*_0_; that is 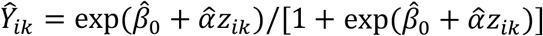 if *Y_ik_* is the disease status (*Y_ik_* = 0,1) for group *k*. Thus, 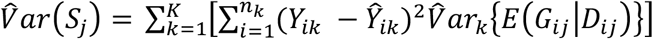. The details of the derivation of the variance of the score test for the multi-group case control set-up as well as for joint rare variant analysis (gene or region based) are provided in the Supplementary document, and in these derivations it is assumed that the covariates are uncorrelated with the genotype, *G_ij_*.

#### Quantitative Traits Analysis

Suppose *Y_i_* is a normally distributed quantitative trait and 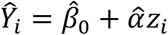 when there is no association. The variance of the score, 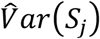, is derived in the Supplementary document. Note that 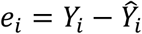 is the residual from the model fitted under *H*_0_ and *Cov*(***e***) = *σ*^2^(I – *H*), where *σ*^2^ is the variance of *Y*, I is an *nxn* identity matrix and *H* is the projection matrix under *H*_0_, i.e. *H* = *Z^t^*(*Z^t^Z*)^−1^*Z^t^* and *Z* is an *n* × (*p* + 1) covariate matrix with 1s in the first column and *p* is the number of covariates. Then 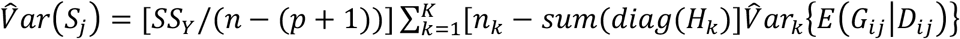 where 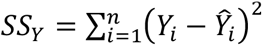, *H_k_* is the section of *H* belonging to group *k*, and *sum*(*diag*(*H_k_*)) is the sum of the diagonals of *H_k_*. This formula can be approximated by a simple form that does not require *H*, i.e. 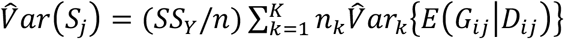. Suppose the NGS data from two groups is combined where the first group was sequenced at HRD and the second at LRD with *n*_1_ < *n*_2_. As opposed to the case-control set up above, the variance of the smaller groups has less weight on the total variance, but the variances of each group are still calculated separately substituting 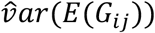 in the variance estimate of the HRD group. The derivation of the variance of the score test for a quantitative trait and for joint rare variant analysis (gene or region based) is provided in the Supplementary document, and again in these derivations it is assumed that the covariates are uncorrelated with the genotype, *G_ij_*. Note that if *Y_i_* is not normally distributed (or more generally, does not follow the distribution assumed), the consistent estimates of the parameters and variance function are not guaranteed (White, 1982), which affects the asymptotic distribution of the score test (Godfrey & Orme, 2001). We recommend that a normal transformation be applied prior to doing association analysis if Y is a highly skewed quantitative phenotype.

For common variant analysis, p-values are calculated using the asymptotic distribution of the score test, 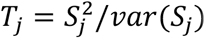, which is distributed as chi-squared with one degree of freedom. For joint rare variant association analysis, VikNGS implements linear and quadratic tests (Derkach, Lawless, & Sun, 2014), with user-defined weights, *w_j_*. With *w_j_*=1, the linear test is analogous to CAST and the quadratic test is analogous to SKAT where 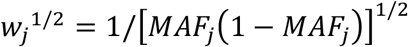 the (Wu et al., 2011). The score statistics for *J* joint SNPs, ***S*** = (*S*_1_, *S*_2_, …, *S*_*J*_) are calculated with the variance of ***S*** estimated by combining the separate covariance matrices for each study stratified by read depth group. Since the distribution of *E*(*G*_*ij*_|*D*_*ij*_) depends on read depth, calculating p-values by permutation is not possible. For binary trait analysis, a bootstrap approach, as in (Derkach, Chiang, et al., 2014) is implemented in VikNGS, where a vector of centered values 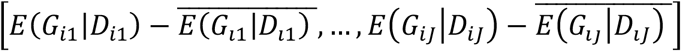 is sampled for each read depth group separately with replacement, e.g. with 10^4^ replicates. In the presence of covariates, the added covariates are also sampled for each read depth group separately with replacement. For quantitative trait analysis, we implement the bootstrap methodology defined in (Lin & Tang, 2011) within each read depth group.

### 2.2 VikNGS package

VikNGS is a C++ cross-platform software package that can run on Windows, Mac and Linux operating systems designed to perform genetic association testing. The package can either be run as a simple command line tool or with a graphical user interface. When run with a user interface, VikNGS also includes options for performing power analysis and interactive data visualizations. To run association tests in VikNGS, a user must provide a multi-sample VCF file and a tab-separated file containing individual-level information as input. As for conventional genetic associations studies, it is important to match all sequenced samples by epidemiologic factors, e.g. ethnicity. Ideally, variants should be called for all samples together to eliminate any systematic bias introduced by different variant calling algorithms. The vRVS methodology can adjust for sequencing parameters, such as sequencing platform, read depth and coverage but other systematic biases in the data can potentially lead to spurious associations. VikNGS enables association testing for both rare and common variants. A BED (browser extensible data) file can be optionally supplied to enable the collapse of variants within genes, exons or any arbitrary interval specified within the file. For computational efficiency, an early stopping procedure is available to terminate the iterative p-value calculations using a method described by (Jiang & Salzman, 2012) if the calculation suggests the p-value is large.

Figure 2 shows a screenshot of the main interface. The user provides input files, filtering parameters and specifies which test statistic to utilize. VikNGS will parse the VCF file, filter variants, potentially collapse variants and will perform association testing all in parallel. For each test performed, VikNGS will produce a p-value and will output summarized variant information and results to a text file. If the graphical user interface is used, an interactive Manhattan-style plot will be produced (Figure 3), and variant-level information can be explored using a tabular view (see Figure S1 and S2).

**Figure 2.**
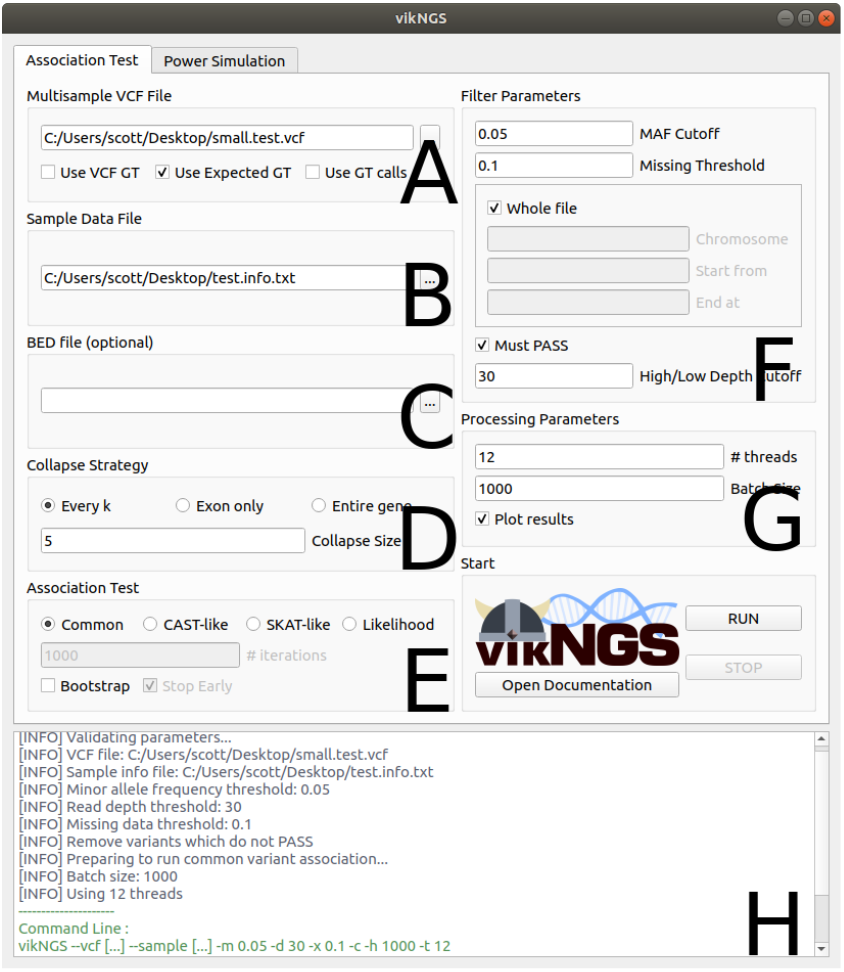
Screenshot of the main VikNGS user interface. **A**: Directory to a multi-sample VCF and how to calculate genotypes from the file. **B**: Directory to a tab-separated file that provides phenotype information, group information and potentially covariates. **C**: Directory to an optional BED format file for variant collapsing. **D**: Collapsing strategy used for rare variant testing. **E**: Details on which association test to run. **F**: parameters used to filter the variants in the file based on minor allele frequency, percentage of data missing or genomic coordinate. **G**: performance settings including how many variants to process in a single thread and the number of threads available. **H**: Output window.

**Figure 3.**
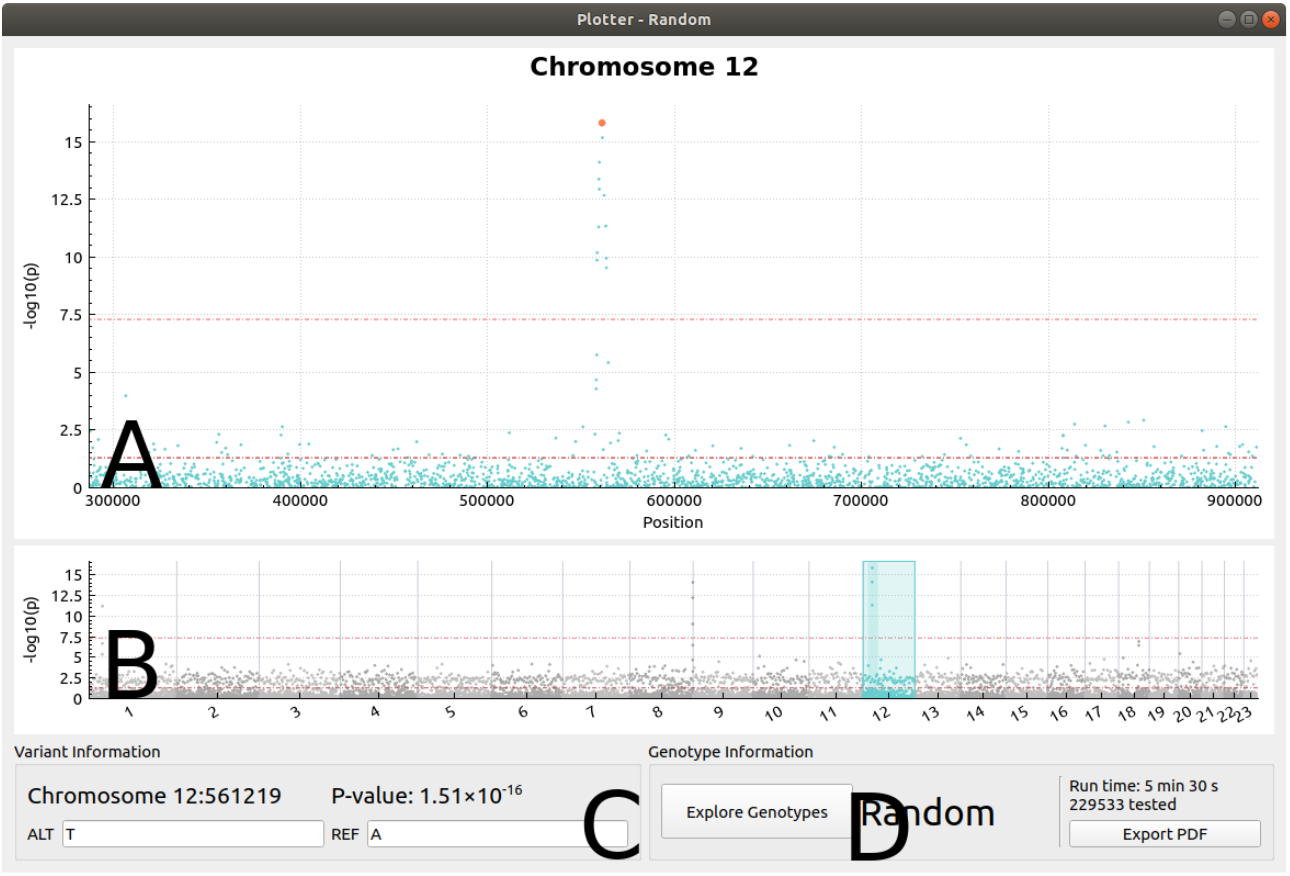
Screenshot of the VikNGS data visualization tool (p-values shown here are randomly generated). **A**: Interactive Manhattan-style plot displaying the association p-values for a single chromosome. **B**: Manhattan plot displaying the entire genome and selector to allow different chromosomes to be explored. **C**: Information displayed when a variant is selected from the chromosome view. **D**: Displays all the variant information and genotypes in a table view (see Figure S1 and S2).

When using the graphical interface, data can be simulated for power and sample size estimation (Figure 4). Parameters such as the number of variants, sample size and effect size are initially specified by a user. VikNGS will first generate a set of true genotypes for each simulated individual, one for each variant. A sequencing experiment is then simulated, producing genotype calls and expected genotypes given the true genotypes and the specified sequencing parameters. A conventional score test using the true genotypes and genotype calls is conducted and the vRVS test is implemented using the expected genotypes. The resulting p-values are visualized. If simulated under the null hypothesis, a Q-Q plot is also displayed to allow comparison of the p-value distribution of the different analytic approaches (Figure 5). A second plot is available to show how power changes as a function of sample size (Figure 6).

**Figure 4.**
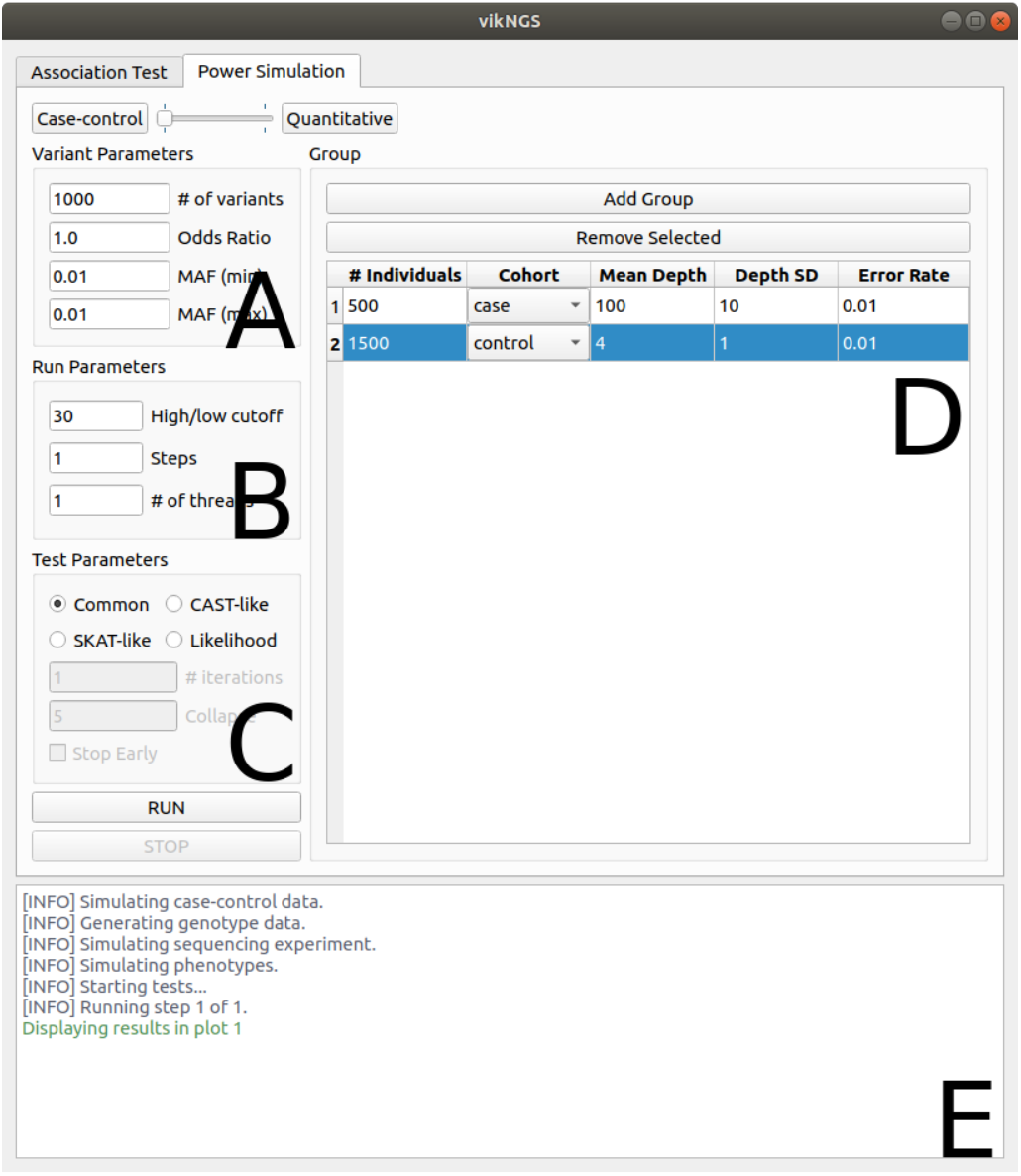
Screenshot of the VikNGS power simulation interface. **A**: Total number of variants to simulate, effect size and a range of minor allele frequency (sampled uniformly at random within the range for each variant). **B**: Defines the depth considered to be high read depth, number of steps with increasing sample size. **C**: Test statistic for p-value calculation and number of threads to use. **D**: Specifies the sequencing simulation parameters and sample size for all groups included in the simulation. **E**: Output window.

**Figure 5.**
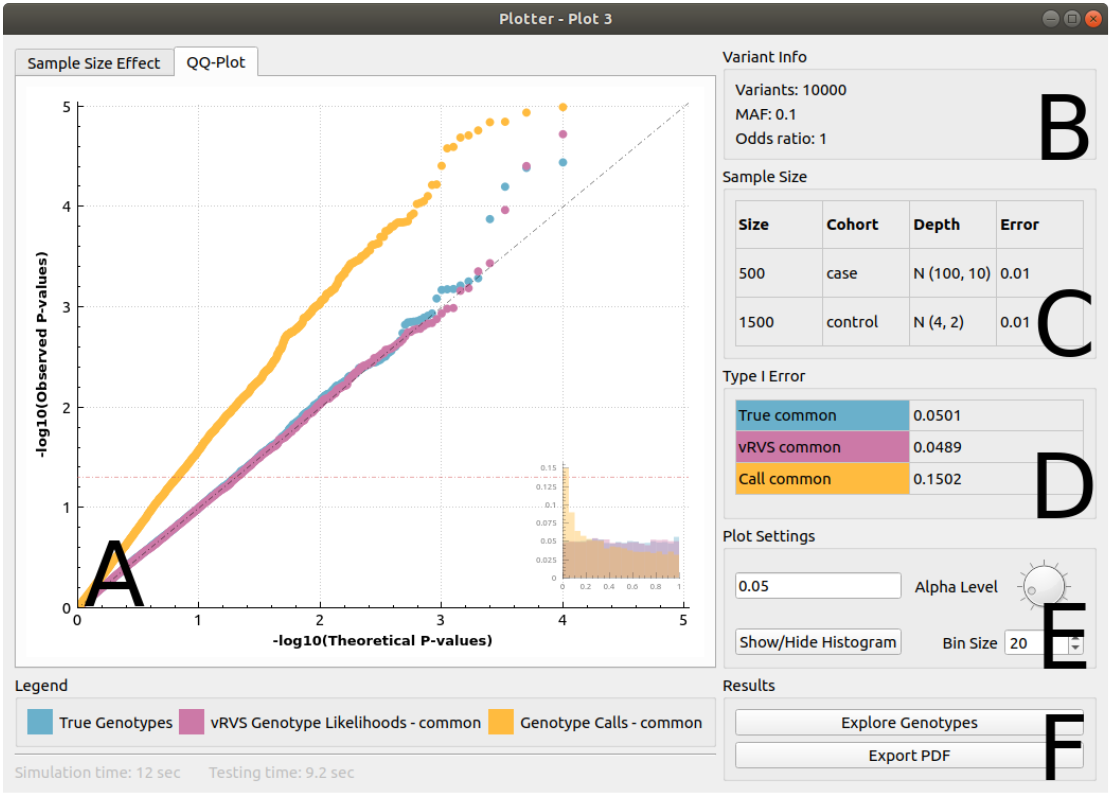
Screenshot of the VikNGS simulated data visualization tool (under null hypothesis). **A**: Q-Q plot and histogram displaying the p-value distribution. **B**: Variant-related parameters in simulation. C: Information regarding which groups were generated as part of the simulation. **D**: View calculated Type I error at a given significance level. **E**: Settings used to interact with the plots (significance level and histogram bin size) **F**: Displays simulated variant information and genotypes in a table view (see Figure S1 and S2), output results to PDF format.

**Figure 6.**
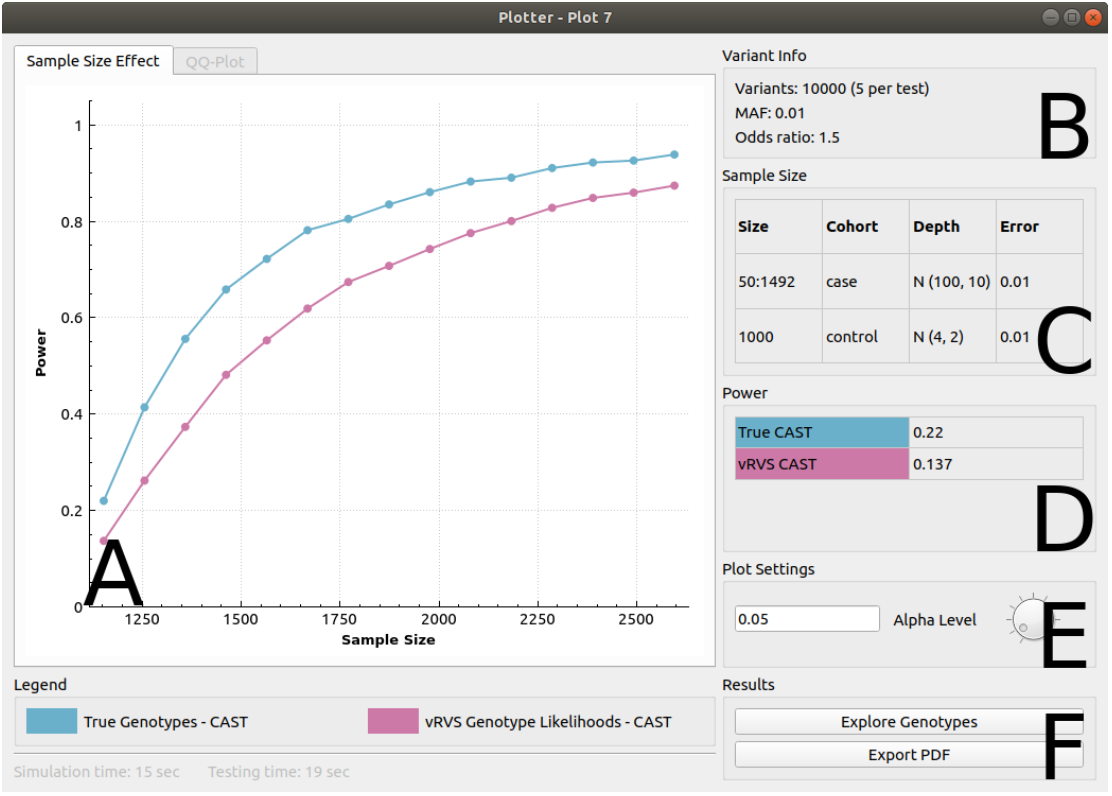
Screenshot of the VikNGS simulated data visualization tool (power with increasing sample size). **A**: Plot displaying the increase of power at a given significance level as total sample size increases. **B**: Variant-related parameters used in simulation. **C**: Information regarding which groups were generated as part of the simulation. **D**: View exact power value error at a given point on the plot in **A** (changes as user interact with the plot). **E**: Significance level used to calculate power. **F**: Displays simulated variant information and genotypes in a table view (see Figure S1 and S2), output results to PDF format.

### 2.3 Simulations

Using the simulation package in VikNGS, we conducted an extensive simulation study to compare the performance of the vRVS to a conventional score test using genotype calls (GC); we used a score test with the true genotypes (true geno) as the gold standard. We investigated the impact of altering sample size, read depth, minor allele frequency (MAF), odds ratio (OR) for case-control simulations and proportion of variation explained by the genetic effect (*R*^2^ - coefficient of determination) for continuous traits. First, true genotypes are generated such that a given individual has either 0, 1 or 2 minor alleles provided by a Binomial distribution with probability of success defined by the MAF. For a case-control study design, genotype data for case and control groups are generated based on the same MAF if under the null hypothesis (OR=1). If the simulation is under the alternative hypothesis, the MAF is simulated to be different between case and control groups, the degree to which they differ being determined by the specified OR. For a quantitative study, once the genotypes are generated the phenotype is simulated based on the normal model, *Y* ~ *N*(*β*_0_ + *β*_1_*g*_*i*_, *σ*^2^). *R*^2^ is converted to *β*_1_ by the equation 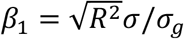 where *σ* is the standard deviation of *Y* and *σ_g_* is the standard deviation of *g*. *β*_0_ is the average of *Y* in the population when there is no genetic or environmental effect.

Given a set of true genotypes, a sequencing experiment is simulated for every individual. For each variant, the read depth is sampled from a Normal distribution with mean and standard deviation specified in the VikNGS interface. The base calling error rate can be specified for each group of simulated individuals. For simulations examined here the error rate is fixed at 0.01, meaning one percent of bases are incorrectly called as one of the three other possible bases. The impact of this base calling error is investigated in the supplementary document (Table S6). After sequence reads are generated, the genotype calls are obtained using the simple Bayesian genotyper (McKenna et al., 2010) which provides the posterior probability of each genotype given the sequence data, and the genotype likelihoods that are used in the vRVS.

The simulation scenarios investigated for case control and quantitative phenotype designs are summarized in Table 1 and Table S1 in the supplementary document, respectively.

**Table 1.**
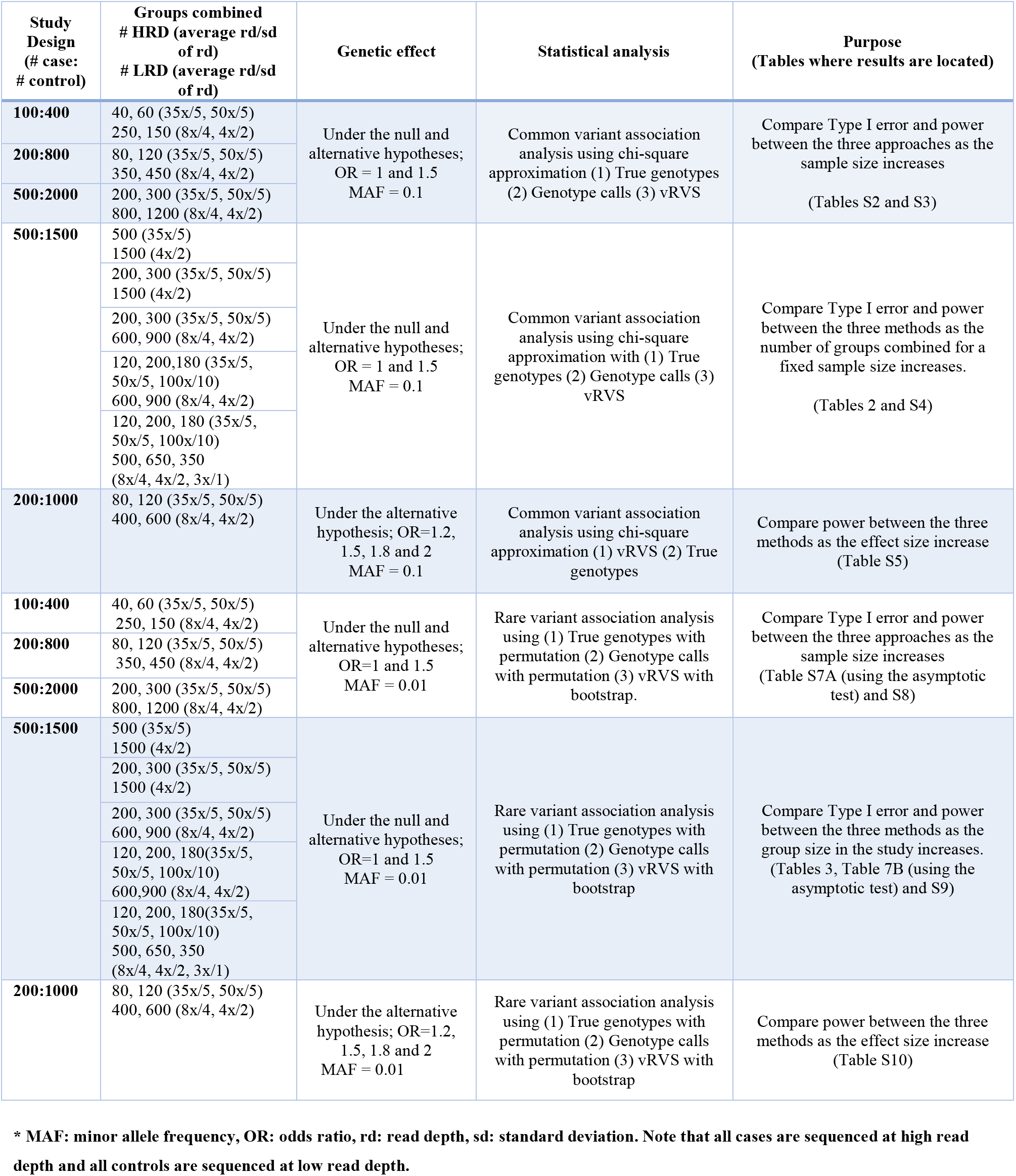
Simulation study for case-control designs

#### 2.3.1 Simulation Results for case-control designs

The empirical Type I error for vRVS, the conventional method using genotype calls and for analysis with the true generating genotypes for common variant analysis are in Table 2 and Table S2 in the supplementary document, respectively. Under all scenarios, Type I error is well controlled for vRVS and is inflated when genotype calls are used for analysis with the score test. With the vRVS the Type I error is controlled regardless of the number of groups combined for a fixed number of cases and controls (Table 2). The empirical Type I error is well controlled for vRVS for different sample sizes (Table S2). In the supplementary document, we see that the power of vRVS increases with sample size and is not affected by the number of groups combined for a fixed sample size (Tables S3 and S4), and that the power of vRVS increases as the effect size gets larger (Table S5), as expected. The power of vRVS is comparable to analyses that use the true genotypes, under all scenarios. Moreover, the empirical Type I error is well controlled with the vRVS when the base calling error is different for cases and controls, while it is inflated when genotypes calls are used for analysis (Table S6).

**Table 2.** Empirical Type I error for common variant analysis with respect to different number of groups combined at the significance level of 0.05 between the vRVS method, a conventional score test using called genotypes (GC) and true generating genotypes (true geno) with 500 HRD cases and 1500 LRD control.

| Number of groups combined | Type of analysis |  |  |
| --- | --- | --- | --- |
|  | True geno | GC | vRVS |
| 2 groups | 0.5020 | 0.1486 | 0.0486 |
| 3 groups | 0.0490 | 0.1536 | 0.0476 |
| 4 groups | 0.0495 | 0.1065 | 0.0478 |
| 5 groups | 0.0506 | 0.1099 | 0.0507 |
| 6 groups | 0.0498 | 0.1423 | 0.0484 |

*Note:* 2 groups indicates 500 cases with 35x and 1500 controls with 4x. 3 groups indicates 200 cases with 35x, 300 cases with 50x and 1500 controls with 4x. 4 groups indicates 200 cases with 35x, 300 cases with 50x and 600 controls with 8x and 900 controls with 4x. 5 groups indicates 120 cases with 35x, 200 cases with 50x, 180 cases with 100x, 600 controls with 8x and 900 controls with 4x. 6 groups indicates 120 cases with 35x, 200 cases with 50x, 180 cases with 100x, 500 controls with 8x, 650 controls with 4x and 350 controls with 3x. Results are based on 10,000 replicates for MAF=0.1. Base calling error is set to 0.01 across all groups. HRD: high read depth, LRD: low read depth, MAF: minor allele frequency.

Simulations for joint rare variant analyses provide similar conclusions. The empirical Type I error with the vRVS is controlled and comparable to an analysis that uses the true genotypes, with CAST being conservative as the number of sequenced groups is varied for rare variant analysis (Table 3). We believe the conservative p-values are due to a low number of possible permutations/bootstrap for low minor allele frequency that do not accurately represent a true null distribution. Empirically, we see that an asymptotic version of CAST using the chi-square approximation results in Type 1 errors that are closer to the nominal value (Tables S7A and Table 7B). The Type I error is inflated in all scenarios when genotype calls are used. The empirical power of vRVS increases with the sample size (Table S8) and effect size (Table S10), is robust to the number of sequenced groups combined (Table S9) and is comparable to an analysis that uses the true genotypes in all scenarios investigated.

**Table 3.** Empirical Type I error for joint rare variant analysis with respect to different number of groups combined at the significance level of 0.05 between the vRVS method, a conventional score test using called genotypes (GC) and true genotype (true geno) with 500 HRD cases and 1500 LRD control.

| Number of groups combined | Type of analysis |  |  |
| --- | --- | --- | --- |
|  | True geno<br>(CAST/SKAT) | GC<br>(CAST/SKAT) | vRVS<br>(CAST/SKAT) |
| 2 groups | 0.0485/0.0525 | 0.2825/0.2030 | 0.0500/0.0455 |
| 3 groups | 0.0390/0.0570 | 0.2805/0.2175 | 0.0415/0.0490 |
| 4 groups | 0.0420/0.0505 | 0.1740/0.1385 | 0.0435/0.0430 |
| 5 groups | 0.0395/0.0505 | 0.1740/0.1330 | 0.0460/0.0525 |
| 6 groups | 0.0420/0.0455 | 0.2120/0.1620 | 0.0410/0.0460 |

*Note:* 2 groups indicates 500 cases with 35x and 1500 controls with 4x. 3 groups indicates 200 cases with 35x, 300 cases with 50x and 1500 controls with 4x. 4 groups indicates 200 cases with 35x, 300 cases with 50x and 600 controls with 8x and 900 controls with 4x. 5 groups indicates 120 cases with 35x, 200 cases with 50x, 180 cases with 100x, 600 controls with 8x and 900 controls with 4x. 6 groups indicates 120 cases with 35x, 200 cases with 50x, 180 cases with 100x, 500 controls with 8x, 650 controls with 4x and 350 controls with 3x. Results are based on 10,000 replicates for MAF=0.01. Base calling error is set to 0.01 across all groups. vRVS is performed through bootstrap (1,000 iterations), GC and true genotype are performed through permutation (1,000 iterations). Each association test used a collapsed group of 5 variants. HRD: high read depth, LRD: low read depth, MAF: minor allele frequency.

#### 2.3.2 Simulation results for quantitative phenotypes

For common and joint rare variant analyses, we choose MAF to be 0.1 (*σ*_*g*_ = 0.42) and 0.01(*σ*_*g*_ = 0.14) respectively. We also let *σ* = 1, *β*_0_ = 0 and *R*^2^ take different values in the simulations. Then the phenotype is generated based on the normal model, *Y* ~ *N*(*β*_0_ + *β*_1_*g*_*i*_, *σ*^2^). We observe that the empirical Type I error for vRVS, the conventional method using genotype calls and for analysis with the true generating genotypes provide similar nominal values for both common and rare variant analysis; that is, the conventional method using genotype calls does not necessarily produce inflated Type I error. As opposed to case control designs, here read depth is not a confounding factor since it is not associated with both genotype calls and the phenotype (Hu et al., 2016). We provide the empirical Type I error for increasing sample size and with different group sizes for common variant association analysis in Tables S11 and S12 and for joint rare variant analysis in Tables S13 and S14. We present the empirical power for common variant analysis with respect to *R*^2^ in Table 4 and for joint rare variant analysis with respect to different number of groups combined in Table 5. As expected, the power of vRVS increases with increasing *R*^2^, is robust to the number of sequenced groups combined and is comparable to an analysis that uses the true genotypes in all scenarios investigated. Moreover, the power of vRVS is always equal or larger than that of the conventional tests based on genotype calls.

**Table 4.** Empirical power for common variant analysis with respect to explained variability (*R*^2^) at the significance level of 0.05 between the vRVS method, a conventional score test using called genotypes (GC) and true genotypes (true geno) for quantitative data analysis with a sample size of 400.

| Explained variance in phenotype ($R^2$) | Type of analysis | | |
| --- | --- | --- | --- |
|  | True geno | GC | vRVS |
| 1% | 0.51061 | 0.45802 | 0.46706 |
| 2% | 0.80261 | 0.74256 | 0.75163 |
| 5% | 0.99165 | 0.98235 | 0.9839 |

*Note:* 4 groups are comprised of 400 samples. 50 samples with 35x, 70 samples with 50x, 200 samples with 4x and 80 samples with 8x. Results are based on 100,000 replicates for MAF=0.1. Base calling error is set to 0.01 across all groups.

**Table 5.** Empirical power for joint rare variant analysis with respect to different number of groups combined at the significance level of 0.05 between the vRVS method, a conventional score test using called genotypes (GC) and true genotypes (true geno) with HRD and LRD groups for quantitative data analysis with sample size of 300.

| Number of groups combined | Type of analysis |  |  |
| --- | --- | --- | --- |
|  | True geno<br>CAST/SKAT | GC<br>CAST/SKAT | vRVS<br>CAST/SKAT |
| 2 groups | 0.9655/0.851 | 0.8995/0.7265 | 0.9195/0.7420 |
| 3 groups | 0.9580/0.8260 | 0.9035/0.7070 | 0.9090/0.7075 |
| 4 groups | 0.9600/0.8470 | 0.9190/0.7445 | 0.9235/0.7555 |
| 5 groups | 0.9550/0.8395 | 0.8990/0.7590 | 0.9100/0.7585 |
| 6 groups | 0.9595/0.84375 | 0.9045/0.7290 | 0.9125/0.7335 |

Note: 2 groups indicates 100 samples with 35x and 200 samples with 4x. 3 groups indicates 50 samples with 35x, 50 samples with 50x and 200 samples with 4x. 4 groups indicates 50 samples with 35x, 50 samples with 50x, 120 samples with 4x and 80 samples with 8x. 5 groups indicates 50 samples with 35x, 30 samples with 50x, 20 samples with 100x, 120 samples with 4x and 80 samples with 8x. 6 groups indicates 50 samples with 35x, 30 samples with 50x, 20 samples with 100x, 90 samples with 4x, 80 samples with 8x and 30 samples with 3x. Results are based on 10,000 replicates for MAF=0.01. Base calling error is set to 0.01 across all groups. vRVS is performed through bootstrap (1,000 iterations), GC and true genotype are performed through permutation (1,000 iterations). Each association test used a collapsed group of 5 variants. $R^2=1\%$ . HRD: high read depth, LRD: low read depth.

#### 2.3.3 Simulations with covariates

It is important for genetic association analyses to adjust for covariates, e.g. sex, age, ethnicity. We extend the functionality of vRVS to enable covariate adjustment, but to do so we break the link between the genotype information and the covariates (see supplementary, variance calculation section). This is equivalent to assuming that the genotype and covariates are not strongly correlated. In this section we investigate through simulation the robustness of vRVS to the correlation between genotype and covariates. We first illustrate the empirical Type I error for a common variant analysis of a binary trait with increasing covariate correlation. Suppose Z is a covariate added to the regression equation, i.e. *p*(*y*_*i*_|*z*_*i*_, *G*_*ij*_) = *exp*(*β*_0_ + *β*_1_*g*_*i*_ + *αz*_*i*_)/(1 + *exp*(*β*_0_ + *β*_1_*g*_*i*_ + *αz*_*i*_)). We investigate the impact of genotype-covariate correlation on Type 1 Error through simulation of 200 cases sequenced at an average of 35x and 600 controls sequenced at an average of 4x with MAF=0.1. Table 6 demonstrates that the Type I error for the vRVS is well controlled for low correlation values (0.01-0.05). As correlation increases (>0.1), the Type I error for vRVS becomes conservative. Results from investigation of smaller (100 cases, 200 controls) and larger sample size (500 case, 1500 control) scenarios are provided in Tables S15 and S16 and demonstrate that, regardless of the sample size, Type I error is controlled in vRVS for low correlations and becomes deflated as correlation increases (>0.1).

**Table 6.** Empirical Type I error for common variant analysis with different correlation values between the genotype, G and the covariate Z at the significance level of 0.05: results provided for the vRVS, a conventional score test using called genotypes (GC) and the true generating genotypes (true geno) with 200 HRD cases and 600 LRD control.

| Cor(G,Z) | Type of analysis |  |  |
| --- | --- | --- | --- |
|  | True geno | GC | vRVS |
| 0 | 0.0499 | 0.0692 | 0.0496 |
| 0.01 | 0.0488 | 0.0687 | 0.0493 |
| 0.02 | 0.4865 | 0.0686 | 0.0484 |
| 0.05 | 0.0500 | 0.0688 | 0.0491 |
| 0.1 | 0.0510 | 0.0706 | 0.0498 |
| 0.2 | 0.0502 | 0.0707 | 0.0449 |
| 0.5 | 0.0503 | 0.0703 | 0.0243 |

*Note:* 200 cases are composed of 80 cases with 35x and 120 cases with 50x and 600 controls are composed of 200 controls with 4x and 400 controls with 8x. Results are based on 100,000 replicates for MAF=0.1. Base calling error is set to 0.01 across all groups. HRD: high read depth, LRD: low read depth. Similar qualitative results observed for quantitative data analysis (Table S19). Likewise, when the covariate is associated with the phenotype Type I error remains controlled with vRVS (Table S20).

#### 2.3.4 Simulations under alternative study designs

Scenarios to this point considered when the HRD case group was smaller than the LRD control group and showed that Type I error is well controlled with the vRVS methodology. If the control group is smaller than the case group, the Type I error has already been shown to be inflated (Table S17) (Derkach, Chiang, et al., 2014). For this experimental design, the estimate of the variance should be based on the full data set, and this option is implemented and must be specified in VikNGS. (Derkach, Chiang, et al., 2014) demonstrated that this approach can cause slight overestimates of the variances but that the Type I error remains controlled.

When both cases and controls are sequenced at high read depth, Type I error (at least for common variant analysis) seems to be controlled on average when using genotype calls (Table S18). However, there will of course be individual sites where read depth differs between the groups and this can likewise translate into spurious findings and power loss (Skotte et al., 2012). For rare variants, even for high read depth sequence, there will be more genotype calling errors and analyses could benefit from methods that use genotype probabilities like the vRVS or the method by Skotte et. al. (Skotte et al., 2012), which will be implemented in VikNGS.

When both cases and controls are sequenced at low read depth, Type I error does not seem to be well controlled for small sample sizes (Table S19). As sample size gets larger (>600), vRVS produces well controlled Type I error while the conventional score test with genotype calls produces inflated Type I error. We also investigated severely unbalanced case-control designs, e.g. the case:control ratio being 50:500 (Table S20). The Type I error rate remains well-controlled using vRVS while the conventional score test with genotype calls produces increasingly inflated Type I error. The power of vRVS remains comparable to that of using the true genotypes (Table S21).

### 2.4 Computational Performance

As an evaluation of the performance capabilities of VikNGS, we used the simulation package to measure the time required to complete a set of association tests. Tests were run on a desktop Ubuntu 18.04 computer with 16 GB of ram and an 8-core AMD Ryzen 7 processor. The wall time used to complete a series of association tests and the number of variants processed per second was computed. The amount of time to process 1 million variants was then inferred. Table 7 shows results for the common variant test. Since the common variant test p-value can be evaluated without bootstrapping/permutation, the test is extremely fast and can be performed on thousands of variants per second. Runtime is dependent on the number of individuals included in the analysis. Modest reductions in runtime can be seen by using multiple threads, with the exception of when the sample size was specified at 500. This is likely because the overhead cost of maintaining 8 threads exceeds the benefits of parallelism when there are a small number of samples.

**Table 7.** Performance of common variant analysis in VikNGS using simulated high read depth data (100x). Samples were equally distributed between cases and controls. 10,000 variants (MAF between 10-30%) were generated for each replicate, each cell contains the average of 3 replicates. All variants were generated under the null hypothesis (odds ratio = 1).

| Test | Sample Size | Variants processed per second (1 thread) | Variants processed per second (8 threads) | 1 million variants (8 threads) |
| --- | --- | --- | --- | --- |
| Common | 500 | $1.2 \times 10^5$ | $7.3 \times 10^4$ | 13 seconds |
| Common | 2000 | $4.0 \times 10^4$ | $5.49 \times 10^4$ | 18 seconds |
| Common | 5000 | $1.7 \times 10^4$ | $3.8 \times 10^4$ | 26 seconds |

Table 8 presents a similar performance analysis for the rare variant tests. Since these tests evaluate a collapsed set of variants and use an iterative p-value calculation, they can be computationally intensive. To reduce this burden, VikNGS includes a method to halt the p-value calculation if enough evidence suggests the test will produce a p-value > 0.05 (Jiang & Salzman, 2012). In VikNGS, this early stopping rule is only applied after a minimum of 10 iterations have completed. The values in Table 8 correspond to running 1,000 iterations which means the smallest p-value that can be resolved is 0.001. For smaller p-values, more iterations must be computed which has a linear impact on the running time. If early stopping is not applied, every set of collapsed variants will take twice as long if the number of iterations is doubled. With early stopping, only collapsed sets with high significance will be affected. In real applications, the vast majority of loci in the genome are expected to be under the null hypothesis of no association, therefore a significant reduction in running time is expected if early stopping is applied.

**Table 8.** Performance of rare variant analysis in VikNGS using simulated high read depth data (100x). Samples were equally distributed between cases and controls. 500 variants (MAF between 1-5%) were generated for each replicate, each cell contains the average of 3 replicates. Each association test used a collapsed group of 5 variants and was run for 1,000 bootstrap/permutation iterations (except early stopping). All variants were generated under the null hypothesis (odds ratio = 1)

| Test | Sample Size | Variants processed per second (1 thread) | Variants processed per second (8 threads) | Variants processed per second (8 threads + early stopping) | 1 million variants (8 threads) | 1 million variants (8 threads + early stopping) |
| --- | --- | --- | --- | --- | --- | --- |
| Rare SKAT | 500 | 48 | 110 | 2419 | 2.5 hours | 6.9 minutes |
| Rare SKAT | 2000 | 18 | 26 | 849 | 10.6 hours | 20.3 minutes |
| Rare SKAT | 5000 | 8 | 10 | 326 | 27.3 hours | 50 minutes |
| Rare CAST | 500 | 83 | 100 | 2557 | 2.8 hours | 6.5 minutes |
| Rare CAST | 2000 | 21 | 25 | 818 | 10.9 hours | 20.3 minutes |
| Rare CAST | 5000 | 8 | 10 | 333 | 27.2 hours | 50 minutes |

Table 7 and Table 8 only consider the time required to run the association test. On a real dataset, data is parsed and processed from a VCF file which requires additional time. Using the same Ubuntu 18.04 desktop as above, the 99 GB VCF file (2,407 individuals) described in Section 3.1 took 12 minutes to parse, filter and run the common variant association test on the 273,241 variants that passed the filtering step. VikNGS is capable of processing data on the scale of the human genome in a reasonable amount of time on a standard desktop computer. Running the command line version on a high-performance computing cluster would yield even shorter runtimes.

## 3 Application to the Genetics of Cystic Fibrosis (CF)

### 3.1 The causal *CFTR* locus

We applied VikNGS to an extremely unbalanced case control study which included NGS data from three independent studies: 101 individuals with CF of European descent sequenced by Complete Genomics at an average read depth of 30x; 1,927 non-CF individuals from the UK10K Consortium (sequenced on Illumina HiSeq 2000, mean read depth of 6.5x) and 379 individuals of European descent from the 1000 Genomes Project Phase 1 (sequenced on a combination of ABI SOLiD and Illumina platforms, mean read depth of 4x). A multi-sample VCF file was generated by merging VCFs across the three datasets using BCFtools since we did not have access to all the BAM files. Ideally the multi-sample VCF should be created by calling variants directly from the BAM sequence files. The resulting 99 GB VCF file contained all variants on chromosome 7. To conduct a common variant analysis, we instructed VikNGS to filter for SNPs with MAF > 5% and remove variants missing more than 10% of data from either cases or controls (these parameters are user defined). 273,241 variants remained after filtering and were tested using the vRVS test statistic with p-values calculated using the asymptotic distribution. The p-values spanning chromosome 7 (Figure 7) demonstrate a strong association at the *CFTR* locus (chr7:117,105,838 - 117,356,025), as expected.

**Figure 7.**
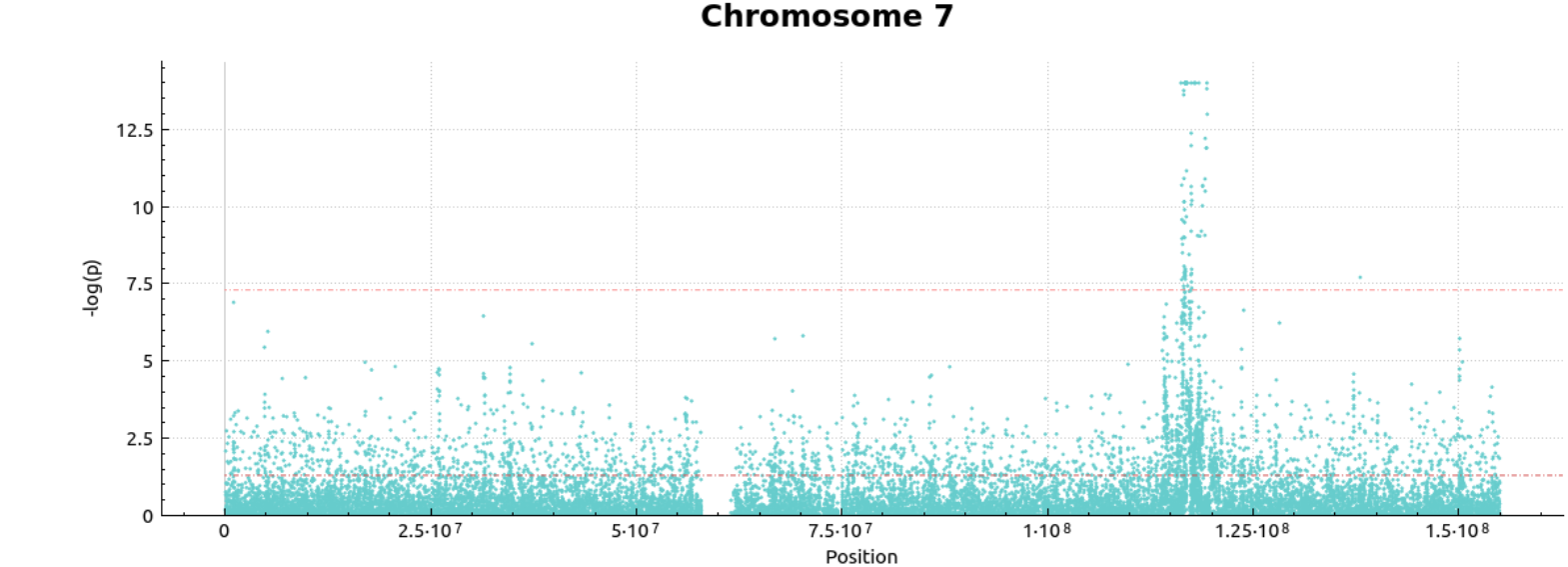
Association test results from an extremely unbalanced design with 101 individuals with cystic fibrosis and 2,306 healthy controls performed by VikNGS. The red dotted lines indicate the p=0.05 and p=5×10^−8^ (conventional genome-wide association significance) levels. The region around CFTR (chr7:117,105,838 - 117,356,025) contains multiple genome-wide significant P-values, with the minimum value VikNGS currently calculates being 1×10^−14^. No other variant across the chromosome exceeded the genome-wide significance threshold.

Although the major CF-causing CF-haplotype displays significant long-range LD, we investigated the distribution of p-values obtained from the *p* arm of chromosome 7, which should be distributed as approximately uniform [0,1] under the null hypothesis although long-range LD with CFTR will impact this slightly. This analysis included 119,955 common SNPs from the start of chromosome 7 to position 58,000,000. We pruned the SNPs using PLINK 1.9 (http://pngu.mgh.harvard.edu/purcell/plink (Purcell et al., 2007), –indep-pairwise 1500 100 0.2) to limit the effect of linkage disequilibrium. 3,845 variants were left after pruning and p-values were computed using vRVS with expected genotypes and with genotype calls for comparison. The distribution of p-values (Figure 8) from the analysis with genotype calls show greater evidence of inflation (λ=1.14) than the vRVS (λ=1.06), consistent with our simulation results and the assumption that the variants on the p arm are not associated with CF.

**Figure 8.**
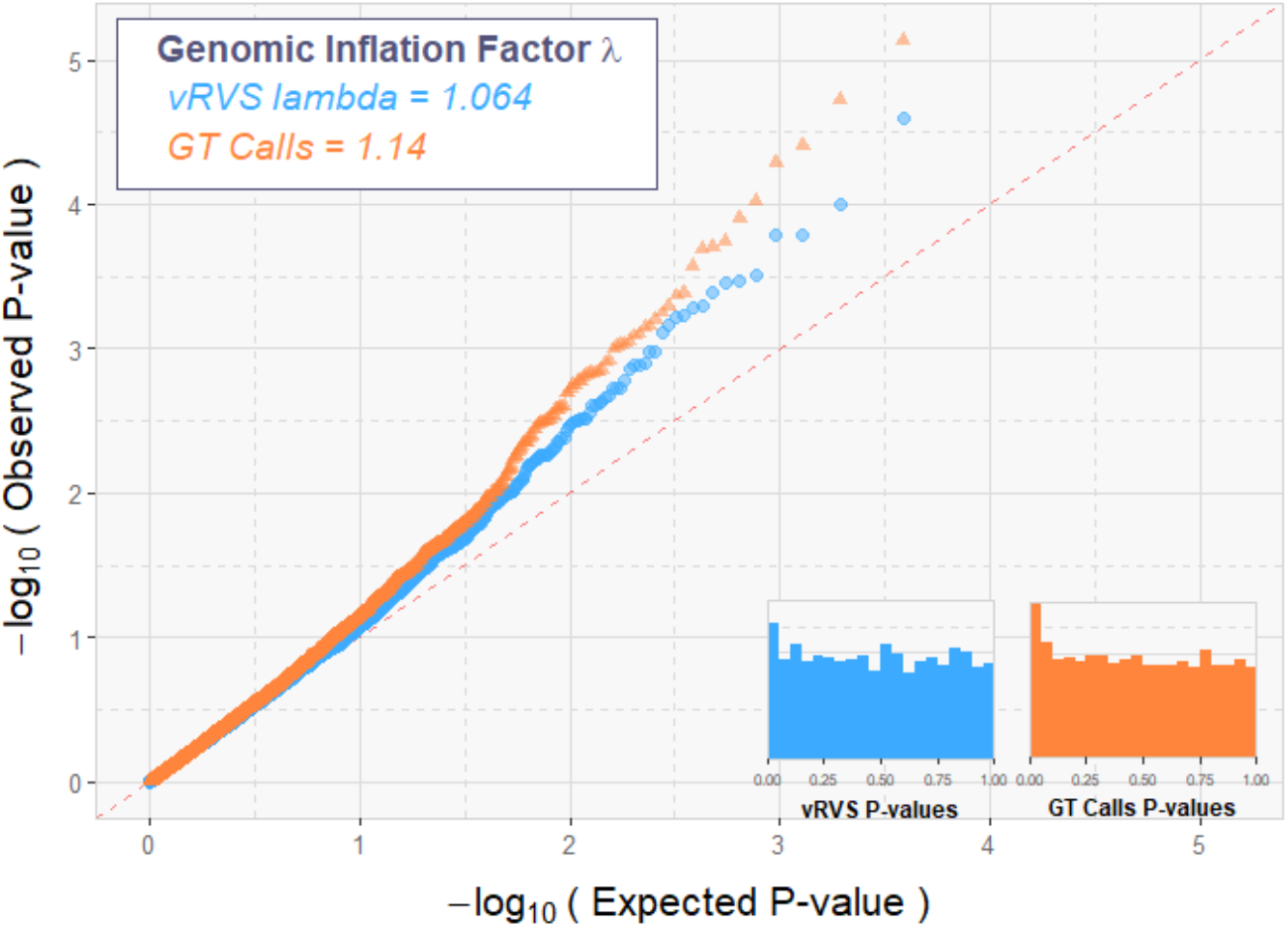
A Q-Q plot was generated from the resulting pruned p-values produced by VikNGS. Orange triangles represent p-values from genotype calls, blue circles correspond to vRVS p-values. P-values derived under the null hypothesis are expected to fall along the dotted diagonal line. A p-value histogram is shown for both distributions and genomic inflation factor λ is provided.

### 3.2 Assessing the contribution of CF modifier gene *SLC26A9* to lung function in the non-CF UK10K population

VikNGS can also implement conventional association testing for NGS data from a single study sequenced using one experimental design. Here we investigate the association evidence between a continuous lung function measurement and variants in a small region of chromosome 1 (chr1:205,860,000 - 205,940,000) near *SLC26A9* that were previously identified to contribute to lung function in older CF (Strug et al., 2016) and non-CF populations (Strug, Stephenson, Panjwani, & Harris, 2018), and hypothesized to influence lung function by improving CFTR function. The association at this locus in non-CF populations is yet to be independently replicated.

In the UK10K data, the common spirometry measures of forced vital capacity (FVC; the total volume of air that can be exhaled from the lungs during a forced expiration) and forced expiratory volume in 1 second (FEV1; the volume exhaled in the first second after maximal inhalation) are available, measured in young participants at the age of 8. A ratio of these two values is a typical measure of lung disease, although the variation in this measure in younger individuals is less pronounced. Quantitative trait association analysis of the FEV1 / FVC ratio at the top CF-associated variant of the *SLC26A9* locus, rs4077468 ((Sun et al., 2012); (Blackman et al., 2013)) in 1,927 non-CF individuals from the UK10K cohort also demonstrated evidence of association using the VikNGS score tests that uses genotype likelihoods (p=0.0392) and genotype calls (p=0.0375).

The association results calculated by VikNGS replicated the other published study in a non-CF population. Publicly available analyses from the UK Biobank (n=307,638) show association with peak expiratory flow and rs4077468 (p=4.35×10^−25^) (McInnes et al., 2018; Strug et al., 2018). The p-values from the VikNGS analysis of the UK10K data reported here are not as small as those previously reported, likely due to the difference in sample size and the limited range of FEV1/FVC in 8 year-olds. However, even with these limitations, this analysis offers further support for variants at this locus contributing to lung function in individuals without CF.

## 1. Discussion

Here we introduce VikNGS, a software application that enables a robust approach for genetic association analysis of common and rare variants from NGS data. Due to sequencing cost considerations, investigators must often limit the sample size of their study population at the cost of power; collaboration across study groups or integration with publicly available data is a pragmatic solution. Integrating data from different sources may introduce systematic biases due to sequencing parameters (error rate, read depth, etc.) which can lead to spurious association findings. Traditional association testing uses called genotypes, the accuracy of which is highly dependent on read depth. Confounding read depth with case-control status will result in an inconsistent distribution of errors in genotype calls. For rare variants, the majority of errors will be mistaken heterozygous calls, resulting in what appears to be an enrichment of minor alleles in one group over another, leading to inflated significance.

VikNGS provides an implementation of the RVS framework (vRVS), built on the concept of the RVS statistic (Derkach, Chiang, et al., 2014) for controlling spurious association due to differences in sequencing parameters. RVS uses expected genotypes instead of genotype calls to enable robust association analysis when combining cases and controls from different experimental designs. We expanded this methodology in VikNGS to enable association analysis on an arbitrary number of datasets for both binary and quantitative trait analysis and allowing for covariate adjustment. We demonstrated through simulation that the vRVS methodology controls Type I error and provides comparable power to analyses if the true genotypes were known, even for severely unbalanced case-control designs. The simulation tool used for evaluating vRVS is included in VikNGS and can be used for power and sample size estimation for study planning. We initially considered several existing simulation software tools to generate sequencing data, e.g. SEQPower (G. T. Wang, Li, Santoz-Cortez, Peng, & Leal, 2014), SimRare (Li, Wang, & Leal, 2010), SeqSIMLA2 (Chung, Tsai, Hsieh, Hung, & Hauser, 2015), but these packages do not offer a simple way to generate and combine data simulated under different sequencing settings.

Genetic association studies that use WGS enable analysis of rare variants which can be analyzed individually or using region-based collapsing association tests, the latter requiring unique statistical methods (Lee, Abecasis, Boehnke, & Lin, 2014). Computational tools have been developed to provide implementations of these methods, eg. EPACTS (http://genome.sph.umich.edu/wiki/EPACTS), PLINK-SEQ (http://atgu.mgh.harvard.edu/plinkseq/), Variant Association Tools (G.T. Wang, Peng, & Leal, 2014), RVTESTS (Zhan, Hu, Li, Abecasis, & Liu, 2016), SCORE-Seq (Lin & Tang, 2011) However, these packages are not designed to analyze data from different sources with varying experimental designs, and they are primarily command line tools rather than implemented through a graphical interface as in VikNGS. To increase the utility of VikNGS, in addition to vRVS we implemented several common and rare variant association analysis approaches using genotype calls.

We have demonstrated the functionality of VikNGS in case-control and quantitative trait settings, using the vRVS framework to combine multiple datasets and for conventional analysis in a single study, respectively. Given 101 individuals with CF and 2,306 non-CF controls, VikNGS detected the *CFTR* locus as associated with CF as expected and was shown to control Type I error better than genotype calls in regions of the chromosome presumed to be approximately under the null hypothesis. Conventional score testing approaches are also implemented in VikNGS for rare and common variant analysis. We implemented this functionality for quantitative trait analysis and replicated an association between lung function and a CF modifier variant in a non-CF population, a gene that has been shown to interact with *CFTR* to improve its function.

The advantage of using sequencing designs over microarray SNP chips in genome-wide association studies is the ability to capture the full allele frequency spectrum enabling rare and common variant analysis. Large sample sizes are required, however, to detect associated rare variants in the data. The vRVS methodology implemented in VikNGS will enable mega-analysis from large consortia despite differences in sequencing design. A current limitation of VikNGS is that the input must be in the format of a multi-sample VCF. This requires a variant calling step to identify polymorphic loci along the genome. As pointed out by (Hu et al., 2016), rare variants can be indistinguishable from base calling errors in low read depth datasets which can lead to truly monomorphic sites being called as rare variants. Including these monomorphic sites in a burden test has been shown to cause inflation in Type I error, even when using vRVS. As this is an issue with variant calling, it is currently outside the scope of VikNGS. However, one can filter out potentially monomorphic sites prior to running a rare variant analysis in VikNGS and using variant calling approaches such as PhredEM (Liao, Satten, & Hu, 2014) which has been shown to significantly reduce the number of monomorphic sites called as polymorphic.

Future developments in VikNGS will allow for more flexibility in input, including the ability to use compressed and indexed VCF files to enable faster parsing and filtering in addition to smaller disk space requirements. Another possible improvement is to allow a set of BAM sequencing files as input to allow greater control over which variants are called and to improve estimates of expected genotypes. Currently, the vRVS framework assumes that specified covariates are uncorrelated with the genotype, *G*_*ij*_. Though weak correlation between the genotype and the covariates has minimal impact, the user should be cognizant of this limitation when specifying covariates. In the current version, vRVS does not accommodate a design where external data is combined with cases and controls who have been sequenced together, although this extension is possible. Lastly the vRVS framework assumes all groups are comparable based on epidemiologic factors, which may be challenging and requires careful consideration. Recently, (Liao, Satten, & Hu, 2018) developed methodology for population structure inference when NGS data is combined across study groups with different experimental designs. Their principal component analysis (PCA) method captures population stratification only, rather than the differences in sequencing properties that would influence traditional PCA approaches. Using covariate adjustment in VikNGS enables inclusion of these PCs for population structure adjustment.

VikNGS is a user-friendly software with a graphical interface that provides a general framework for simulation and genetic association using NGS combined across different study cohorts. The software is fast and user friendly, offering efficient parallel computations that can be run from the command line or using a visual interface on Windows, Linux or Mac operating systems. VikNGS is freely available at http://www.tcag.ca/tools/index.html. Detailed documentation is available at https://VikNGSdocs.readthedocs.io/en/latest/.

