## Supplementary Data for "VikNGS: A C++ Variant Integration Kit for Next Generation Sequencing association analysis"

### 1. VikNGS Interface

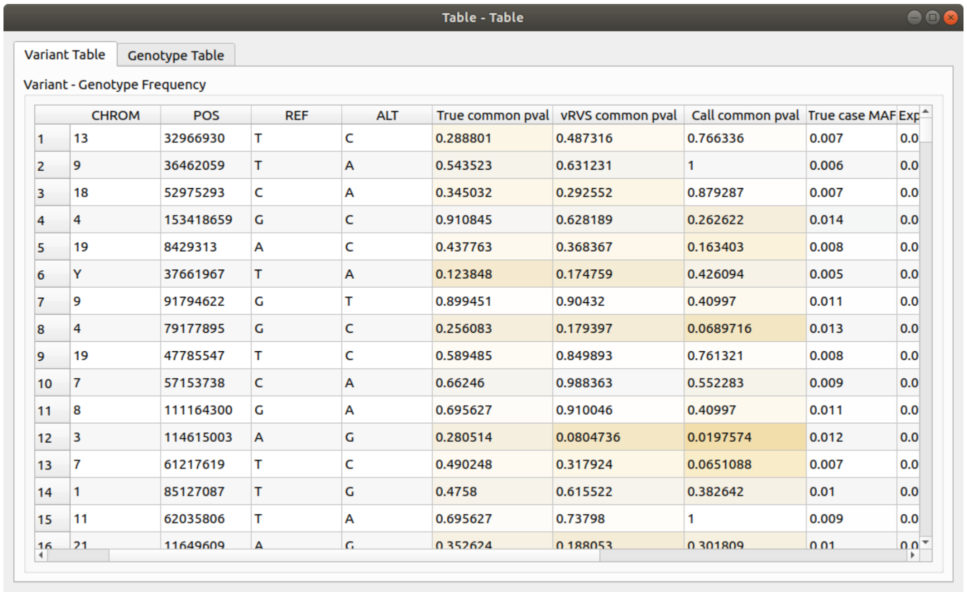
Table - Table

Variant Table | Genotype Table

Variant - Genotype Frequency

| | CHROM | POS | REF | ALT | True common pval | vRVS common pval | Call common pval | True case MAF | Exp |
|---|---|---|---|---|---|---|---|---|---|
| 1 | 13 | 32966930 | T | C | 0.288801 | 0.487316 | 0.766336 | 0.007 | 0.0 |
| 2 | 9 | 36462059 | T | A | 0.543523 | 0.631231 | 1 | 0.006 | 0.0 |
| 3 | 18 | 52975293 | C | A | 0.345032 | 0.292552 | 0.879287 | 0.007 | 0.0 |
| 4 | 4 | 153418659 | G | C | 0.910845 | 0.628189 | 0.262622 | 0.014 | 0.0 |
| 5 | 19 | 8429313 | A | C | 0.437763 | 0.368367 | 0.163403 | 0.008 | 0.0 |
| 6 | Y | 37661967 | T | A | 0.123848 | 0.174759 | 0.426094 | 0.005 | 0.0 |
| 7 | 9 | 91794622 | G | T | 0.899451 | 0.90432 | 0.40997 | 0.011 | 0.0 |
| 8 | 4 | 79177895 | G | C | 0.256083 | 0.179397 | 0.0689716 | 0.013 | 0.0 |
| 9 | 19 | 47785547 | T | C | 0.589485 | 0.849893 | 0.761321 | 0.008 | 0.0 |
| 10 | 7 | 57153738 | C | A | 0.66246 | 0.988363 | 0.552283 | 0.009 | 0.0 |
| 11 | 8 | 111164300 | G | A | 0.695627 | 0.910046 | 0.40997 | 0.011 | 0.0 |
| 12 | 3 | 114615003 | A | G | 0.280514 | 0.0804736 | 0.0197574 | 0.012 | 0.0 |
| 13 | 7 | 61217619 | T | C | 0.490248 | 0.317924 | 0.0651088 | 0.007 | 0.0 |
| 14 | 1 | 85127087 | T | G | 0.4758 | 0.615522 | 0.382642 | 0.01 | 0.0 |
| 15 | 11 | 62035806 | T | A | 0.695627 | 0.73798 | 1 | 0.009 | 0.0 |
| 16 | 21 | 11649609 | A | G | 0.352624 | 0.188053 | 0.301809 | 0.01 | 0.0 |

Figure S1. Screenshot of table containing simulated variant information displayed by VikNGS. A similar table can be viewed if vikNGS is run with real data. This table enables exploration of p-values and minor allele frequency (MAF) of the variants generated and tested in VikNGS. Selecting a particular row in the table allows exploration of the genotypes of that variant in the "Genotype Table" tab (see Figure S2).

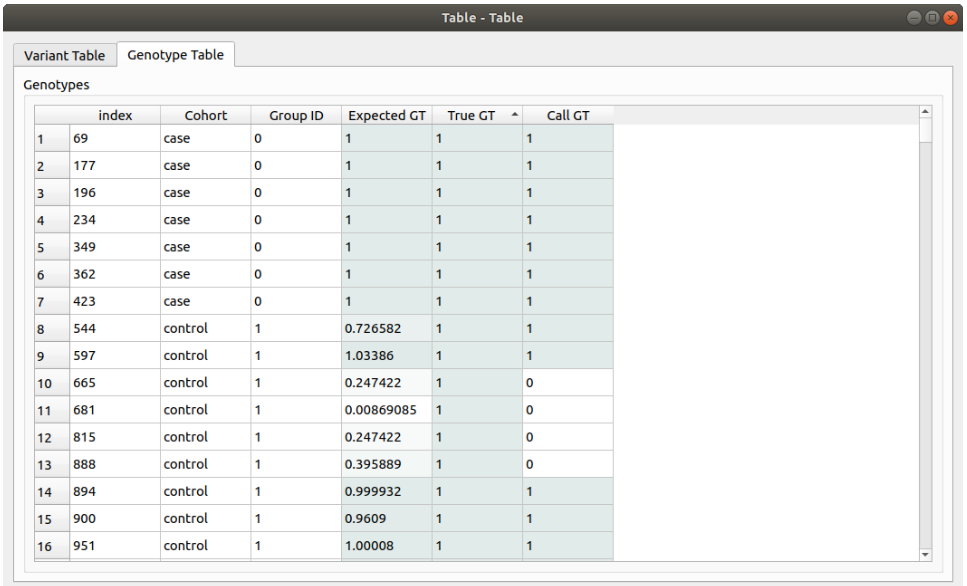
Table - Table

Variant Table | Genotype Table

Genotypes

| | index | Cohort | Group ID | Expected GT | True GT | Call GT |
|---|---|---|---|---|---|---|
| 1 | 69 | case | 0 | 1 | 1 | 1 |
| 2 | 177 | case | 0 | 1 | 1 | 1 |
| 3 | 196 | case | 0 | 1 | 1 | 1 |
| 4 | 234 | case | 0 | 1 | 1 | 1 |
| 5 | 349 | case | 0 | 1 | 1 | 1 |
| 6 | 362 | case | 0 | 1 | 1 | 1 |
| 7 | 423 | case | 0 | 1 | 1 | 1 |
| 8 | 544 | control | 1 | 0.726582 | 1 | 1 |
| 9 | 597 | control | 1 | 1.03386 | 1 | 1 |
| 10 | 665 | control | 1 | 0.247422 | 1 | 0 |
| 11 | 681 | control | 1 | 0.00869085 | 1 | 0 |
| 12 | 815 | control | 1 | 0.247422 | 1 | 0 |
| 13 | 888 | control | 1 | 0.395889 | 1 | 0 |
| 14 | 894 | control | 1 | 0.999932 | 1 | 1 |
| 15 | 900 | control | 1 | 0.9609 | 1 | 1 |
| 16 | 951 | control | 1 | 1.00008 | 1 | 1 |

Figure S2. Screenshot of table containing simulated genotype information displayed by VikNGS. A similar table can be viewed if vikNGS is run with real data. This table enables comparison of called, expected and true genotypes of a single variant generated and tested in VikNGS. Using real data, true genotype values are not known and the "True GT" column is not shown.

#### 2. Simulations

Table S1. Simulation study plan for quantitative phenotype

| Sample size | Groups combined<br>nHRD (average/sd of read depth)<br>nLRD (average/sd of read depth) | Data generated | Genetic effect | Genetic effect | Purpose<br>(Table where results are located) |
|---|---|---|---|---|---|
| 100 | 20, 40 (35x/5, 50x/5)<br>25, 15 (8x/4, 4x/2) | 100,000 replicates for MAF=0.1 | Under the null hypothesis; $R^2$=0% | Common variant association analysis using chi-square approximation (1)True genotypes (2) Genotype calls (3) vRVS | Compare Type I error between the three approaches as the sample size increases<br><br>(Table S11) |
| 500 | 100, 120 (35x/5, 50x/5)<br>200, 80 (8x/4, 4x/2) | | | | |
| 1000 | 100, 250 (35x/5, 50x/5)<br>450, 200 (4x/2, 8x/4) | | | | |
| 300 | 100 (35x/5)<br>200 (4x/2)<br><br>50,50 (35x/5, 50x/5)<br>200 (4x/2)<br><br>50,50 (35x/5, 50x/5)<br>120, 80 (4x/2, 8x/4)<br><br>50, 30, 50 (35x/5, 50x/5, 100x/10)<br>100, 80 (4x/2, 8x/4)<br><br>50, 30 , 20(35x/5, 50x/5, 100x/10)<br>90, 80, 30 (4x/2, 8x/4, 3x/1) | 100,000 replicates for MAF=0.1 | Under the null and alternative hypotheses; $R^2$=0% | Common variant association analysis using chi-square approximation (1) True genotypes (2) Genotype calls (3) vRVS | Compare Type I error between the three methods as the number of groups combined for a fixed sample size increases.<br><br>(Table S12) |
| 400 | 50, 70 (35x/5, 50x/5)<br>200, 80 (4x/2, 8x/4) | 100,000 replicates for MAF=0.1 | Under the alternative hypothesis $R^2$=1%<br>Under the alternative hypothesis $R^2$=2%<br>Under the alternative hypothesis $R^2$=5% | Common variant association analysis using chi-square approximation (1) True genotypes (2) Genotype calls (3) vRVS | Compare power between the three methods as the effect size increase (Table 4) |
| 100 | 20, 40 (35x/5, 50x/5)<br>25, 15 (8x/4, 4x/2) | 10,000 replicates for MAF=0.01 | Under the null and alternative hypotheses; $R^2$=0% | Rare variant association analysis using (1) True genotypes with permutation (2) Genotype calls with permutation (3) vRVS with bootstrap | Compare Type I error between the three approaches as the sample size increases (Table S13) |
| 500 | 100, 120 (35x/5, 50x/5)<br>200, 80 (8x/4, 4x/2) | | | | |
| 1000 | 100, 250 (35x/5, 50x/5)<br>450, 200 (4x/2, 8x/4) | | | | |
| 500 | 100 (35x/5)<br>200 (4x/2)<br><br>50,50 (35x/5, 50x/5)<br>200 (4x/2)<br><br>50,50 (35x/5, 50x/5)<br>120, 80 (4x/2, 8x/4)<br><br>50, 30, 50 (35x/5, 50x/5, 100x/10)<br>100, 80 (4x/2, 8x/4)<br><br>50, 30 , 20(35x/5, 50x/5, 100x/10)<br>90, 80, 30 (4x/2, 8x/4, 3x/1) | 10,000 replicates for MAF=0.01 | Under the null and alternative hypotheses; $R^2$=0% and $R^2$==1% | Rare variant association analysis using (1) True genotypes with permutation (2) Genotype calls with permutation (3) vRVS with bootstrap | Compare Type I error and power between the three methods as the group size in the study increases. (Table 5 and Table S14) |

*nHRD: number of samples sequenced at high read depth, nLRD: number of samples sequenced at low read depth, MAF: minor allele frequency. $R^2$ is the variability in the quantitative trait explained by the SNP, under the null hypothesis, it is taken 0. The standard deviations of the SNP generated and quantitative phenotype are indicated as $\sigma_x$ and $\sigma_Y$ respectively in the table. Based on $R^2$, $\sigma_x$ and $\sigma_Y$, $\beta_1$ is calculated and the quantitative phenotype $Y$ is generated based on linear regression model, i.e. Y~$\beta_0 + \beta_1 SNP$. For the simulation study in common variant analyses, we take $\beta_0$=0, $\sigma_Y = 1$, $\sigma_x = \sqrt{2MAF(1-MAF)}$.

#### 2.1. Simulation Results for case-control designs

Table S2. Empirical Type I error for common variant analysis with respect to sample size at the significant level of 0.05 between the vRVS method, a conventional score test using called genotypes (GC) and true genotype (true geno) with HRD cases and LRD control.

| Sample size (case:control) | Type of analysis | | |
|---|---|---|---|
| | True geno | GC | vRVS |
| 100:400 | 0.04905 | 0.06336 | 0.04908 |
| 200:800 | 0.04925 | 0.07775 | 0.04914 |
| 500:2000 | 0.04951 | 0.11579 | 0.04883 |

*Note:* 100 cases are composed of two groups (40 cases with 35x and 60 cases with 50x) and 400 controls are composed of two groups (250 controls with 8x and 150 controls with 4x). 200 cases are composed of two groups (80 cases with 35x and 120 cases with 50x) and 800 controls are composed of two groups (350 controls with 8x and 450 controls with 4x). 500 cases are composed of two groups (200 cases with 35x and 300 cases with 50x) and 2000 controls are composed of two groups (800 controls with 8x and 1200 controls with 4x). Results are based on 100,000 replicates for MAF=0.1. Base calling error is set to 0.01 across all groups. HRD: high read depth, LRD:low read depth, MAF=minor allele frequency.

Table S3. Empirical power for common variant analysis with respect to sample size at the significant level of 0.05 between the vRVS method and the score test with true genotype (true geno) with HRD cases and LRD control.

| Number of groups combined | Type of analysis | |
|---|---|---|
| | True geno | vRVS |
| 100:400 | 0.4118 | 0.4041 |
| 200:800 | 0.6718 | 0.6548 |
| 500:2000 | 0.9635 | 0.9577 |

*Note:* 100 cases are composed of two groups (40 cases with 35x and 60 cases with 50x) and 400 controls are composed of two groups (250 controls with 8x and 150 controls with 4x). 200 cases are composed of two groups (80 cases with 35x and 120 cases with 50x) and 800 controls are composed of two groups (350 controls with 8x and 450 controls with 4x). 500 cases are composed of two groups (200 cases with 35x and 500 cases with 50x) and 2000 controls are composed of two groups (800 controls with 8x and 1200 controls with 4x). Results are based on 10,000 replicates for MAF=0.1. Base calling error is set to 0.01 across all groups. OR=1.5. HRD: high read depth, LRD:low read depth, MAF=minor allele frequency, OR:odds ratio.

Table S4. Empirical power for common variant analysis with respect to different number of groups combined at the significant level of 0.05 between the vRVS method and the score test using called true genotype (true geno) with 500 HRD cases and 1500 LRD control.

| Number of groups combined | Type of analysis | |
|---|---|---|
| | True geno | vRVS |
| 2 groups | 0.9512 | 0.9376 |
| 3 groups | 0.9533 | 0.9397 |
| 4 groups | 0.9517 | 0.9424 |
| 5 groups | 0.9499 | 0.9400 |
| 6 groups | 0.9515 | 0.9406 |

*Note:* 2 groups indicates 500 cases with 35x and 1500 controls with 4x. 3 groups indicates 200 cases with 35x, 300 cases with 50x and 1500 controls with 4x. 4 groups indicates 200 cases with 35x, 300 cases with 50x and 600 controls with 8x and 900 controls with 4x. 5 groups indicates 120 cases with 35x, 200 cases with 50x, 180 cases with 100x, 600 controls with 8x and 900 controls with 4x. 6 groups indicates 120 cases with 35x, 200 cases with 50x, 180 cases with 100x, 500 controls with 8x, 650 controls with 4x and 350 controls with 3x. Results are based on 10,000 replicates for MAF =0.1. Base calling error is set to 0.01 across all groups. OR equals to 1.5.

Table S5. Empirical power for common variant analysis with respect to effect size at the significant level of 0.05 between the vRVS method and the score test using true genotype (true geno) with 200 HRD cases and 1000 LRD control.

| Effect size (OR) | Type of analysis | |
|---|---|---|
| | True geno | vRVS |
| 1.2 | 0.1925 | 0.1871 |
| 1.5 | 0.6898 | 0.6737 |
| 1.8 | 0.9568 | 0.9495 |
| 2.0 | 0.9903 | 0.9892 |

*Note:* 200 cases are composed of two groups (80 cases with 35x and 120 cases with 50x) and 1000 controls are composed of two groups (400 controls with 8x and 600 controls with 4x). Results are based on 10,000 replicates for MAF=0.1. Base calling error is set to 0.01 across all groups.

Table S6. Empirical Type I error for common variant analysis with respect to different error rates in control groups at the significant level of 0.05 between the vRVS method, a conventional score test using called genotypes (GC) and true genotype (true geno) with HRD cases and LRD control.

| Sample size (case:control) | Type of analysis | | |
|---|---|---|---|
| | Error rate | GC | vRVS |
| 100:400 | 0.005 | 0.06043 | 0.05093 |
| | 0.015 | 0.05953 | 0.05098 |
| | 0.020 | 0.05600 | 0.05127 |
| | 0.030 | 0.05717 | 0.05094 |
| 200:800 | 0.005 | 0.09199 | 0.04995 |
| | 0.015 | 0.07577 | 0.04923 |
| | 0.020 | 0.07389 | 0.04977 |
| | 0.030 | 0.08211 | 0.05004 |

*Note:* 100 cases are composed of two groups (40 cases with 35x and 60 cases with 50x) and 400 controls are composed of two groups (250 controls with 8x and 150 controls with 4x). 200 cases are composed of two groups (80 cases with 35x and 120 cases with 50x) and 800 controls are composed of two groups (350 controls with 8x and 450 controls with 4x). Results are based on 100,000 replicates for MAF=0.1. Base calling error for controls are set to 0.005, 0.015, 0.020 and 0.030 while it is set to 0.01 for cases in each scenario.

Table S7A. Empirical Type I error for joint rare variant analysis with respect to sample size at the significant level of 0.05 between the vRVS method, a conventional score test using called genotypes (GC) and true genotype (true geno) with HRD cases and LRD control.

| Sample size (case:control) | Type of analysis | | |
|---|---|---|---|
| | True geno CAST | GC CAST | vRVS CAST |
| 100:400 | 0.0477 | 0.0720 | 0.0459 |
| 200:800 | 0.0488 | 0.1111 | 0.0474 |
| 500:2000 | 0.0489 | 0.2117 | 0.0478 |

*Note:* 100 cases are composed of two groups (40 cases with 35x and 60 cases with 50x) and 400 controls are composed of two groups (250 controls with 8x and 150 controls with 4x). 200 cases are composed of two groups (80 cases with 35x and 120 cases with 50x) and 800 controls are composed of two groups (350 controls with 8x and 450 controls with 4x). 500 cases are composed of two groups (200 cases with 35x and 300 cases with 50x) and 2000 controls are composed of two groups (800 controls with 8x and 1200 controls with 4x). Results are based on 100,000 replicates for MAF=0.01. Base calling error is set to 0.01 across all groups. Each association test used a collapsed group of 5 variants, and is performed through **asymptotic chi-square approach** with 1 degrees of freedom.

Table 7B. Empirical Type I error for joint rare variant analysis with respect to different number of groups combined at the significance level of 0.05 between the vRVS method, a conventional score test using called genotypes (GC) and true genotype (true geno) with 500 HRD cases and 1500 LRD control.

| Number of groups combined | Type of analysis | | |
|---|---|---|---|
| | True geno CAST | GC CAST | vRVS CAST |
| 2 groups | 0.0500 | 0.3040 | 0.0540 |
| 3 groups | 0.0465 | 0.3045 | 0.0410 |
| 4 groups | 0.0520 | 0.1895 | 0.0520 |
| 5 groups | 0.0530 | 0.1825 | 0.0505 |
| 6 groups | 0.0485 | 0.2235 | 0.0445 |

*Note:* 2 groups indicates 500 cases with 35x and 1500 controls with 4x. 3 groups indicates 200 cases with 35x, 300 cases with 50x and 1500 controls with 4x. 4 groups indicates 200 cases with 35x, 300 cases with 50x and 600 controls with 8x and 900 controls with 4x. 5 groups indicates 120 cases with 35x, 200 cases with 50x, 180 cases with 100x, 600 controls with 8x and 900 controls with 4x. 6 groups indicates 120 cases with 35x, 200 cases with 50x, 180 cases with 100x, 500 controls with 8x, 650 controls with 4x and 350 controls with 3x. Results are based on 10,000 replicates for MAF=0.01. Base calling error is set to 0.01 across all groups. Each association test used a collapsed group of 5 variants, and is performed through **asymptotic chi-square approach** with 1 degrees of freedom. LRD: low read depth, MAF: minor allele frequency.

Table S8. Empirical power for joint rare variant analysis with respect to sample size at the significant level of 0.05 between the vRVS method and the conventional score test using true genotype (true geno) with HRD cases and LRD control.

| Sample size (case:control) | Type of analysis | |
|---|---|---|
| | True geno (CAST/SKAT) | vRVS (CAST/SKAT) |
| 100:400 | 0.5220/0.4000 | 0.4930/0.3745 |
| 200:800 | 0.7820/0.6620 | 0.7665/0.6360 |
| 500:2000 | 0.9845/0.9500 | 0.9820/0.9405 |

*Note:* 100 cases are composed of two groups (40 cases with 35x and 60 cases with 50x) and 400 controls are composed of two groups (250 controls with 8x and 150 controls with 4x). 200 cases are composed of two groups (80 cases with 35x and 120 cases with 50x) and 800 controls are composed of two groups (350 controls with 8x and 450 controls with 4x). 500 cases are composed of two groups (200 cases with 35x and 300 cases with 50x) and 2000 controls are composed of two groups (800 controls with 8x and 1200 controls with 4x). Results are based on 10,000 replicates for MAF=0.01. Base calling error is set to 0.01 across all groups. vvRVS is performed through bootstrap (1,000 iterations), true genotype is performed through permutation (1,000 iterations). Each association test used a collapsed group of 5 variants. Odds ratio = 1.5.

Table S9. Empirical power for joint rare variant analysis with respect to different number of groups combined at the significant level of 0.05 between the vRVS method and the conventional score test using true genotype (true geno) with 500 HRD cases and 1500 LRD control.

| Number of groups combined | Type of analysis | |
|---|---|---|
| | True geno CAST/SKAT | vRVS CAST/SKAT |
| 2 groups | 0.7725/0.5975 | 0.7385/0.5310 |
| 3 groups | 0.7705/0.6150 | 0.7180/0.5540 |
| 4 groups | 0.7700/0.6135 | 0.7350/0.5645 |
| 5 groups | 0.7755/0.6120 | 0.7470/0.5625 |
| 6 groups | 0.7835/0.6115 | 0.7525/0.5625 |

*Note:* 2 groups indicates 500 cases with 35x and 1500 controls with 4x. 3 groups indicates 200 cases with 35x, 300 cases with 50x and 1500 controls with 4x. 4 groups indicates 200 cases with 35x, 300 cases with 50x and 600 controls with 8x and 900 controls with 4x. 5 groups indicates 120 cases with 35x, 200 cases with 50x, 180 cases with 100x, 600 controls with 8x and 900 controls with 4x. 6 groups indicates 120 cases with 35x, 200 cases with 50x, 180 cases with 100x, 500 controls with 8x, 650 controls with 4x and 350 controls with 3x. Results are based on 10,000 replicates for MAF=0.01. Base calling error is set to 0.01 across all groups. vvRVS is performed through bootstrap (1,000 iterations), true genotype is performed through permutation (1,000 iterations). Each association test used a collapsed group of 5 variants. Odds ratio = 1.5.

Table S10. Empirical power for joint rare variant analysis with respect to effect size at the significant level of 0.05 between the vRVS method and the conventional score test using true genotype (true geno) with 200 HRD cases and 1000 LRD control.

| Effect size (OR) | Type of analysis | |
|---|---|---|
| | True geno CAST/SKAT | vRVS CAST/SKAT |
| 1.2 | 0.2600/0.1805 | 0.2380/0.1790 |
| 1.5 | 0.7990/0.6830 | 0.7805/0.6550 |
| 2.0 | 0.9970/0.9910 | 0.9965/0.9875 |

*Note:* 200 cases are composed of two groups (80 cases with 35x and 120 cases with 50x) and 1000 controls are composed of two groups (400 controls with 8x and 600 controls with 4x). RResults are based on 10,000 replicates for MAF=0.01. Base calling error is set to 0.01 across all groups. vvRVS is performed through bootstrap (1,000 iterations), true genotype is performed through permutation (1,000 iterations). Each association test used a collapsed group of 5 variants.

#### 2.2. Simulation results for quantitative phenotypes

Table S11. Empirical Type I error for common variant analysis with respect to sample size at the significant level of 0.05 between the vRVS method, a conventional score test using called genotypes (GC) and true genotype (true geno) with HRD and LRD groups for quantitative data analysis.

| Sample size | Type of analysis | | |
|---|---|---|---|
| | True geno | GC | vRVS |
| 100 | 0.0503 | 0.0504 | 0.0507 |
| 500 | 0.0507 | 0.0504 | 0.0501 |
| 1000 | 0.0497 | 0.0506 | 0.0507 |

*Note:* 100 samples are composed of 4 groups; 20 samples with 35x, 40 samples with 50x, 25 samples with 4x and 15 samples with 8x. 500 samples are composed of 4 groups; 100 samples with 35x, 120 samples with 50x, 200 samples with 4x and 80 samples with 8x. 1000 samples are composed of 4 groups; 100 samples with 35x, 250 samples with 50x, 450 samples with 4x and 200 samples with 8x. Results are based on 100,000 replicates for MAF=0.1. Base calling error is set to 0.01 across all groups. HRD: high read depth, LRD: low read depth.

Table S12. Empirical Type I error for common variant analysis with respect to different number of groups combined at the significant level of 0.05 between the vRVS method, a conventional score test using called genotypes (GC) and true genotype (true geno) with HRD and LRD groups for quantitative data analysis.

| Number of groups combined | Type of analysis | | |
|---|---|---|---|
| | True geno | GC | vRVS |
| 2 groups | 0.0500 | 0.0506 | 0.0507 |
| 3 groups | 0.0491 | 0.0491 | 0.0495 |
| 4 groups | 0.0501 | 0.0500 | 0.0504 |
| 5 groups | 0.0497 | 0.0496 | 0.0496 |
| 6 groups | 0.0494 | 0.0493 | 0.0495 |

*Note:* 2 groups indicates 100 samples with 35x and 200 samples with 4x. 3 groups indicates 50 samples with 35x, 50 samples with 50x and 200 samples with 4x. 4 groups indicates 50 samples with 35x, 50 samples with 50x, 120 samples with 4x and 80 samples with 8x. 5 groups indicates 50 samples with 35x, 30 samples with 50x, 20 samples with 100x, 120 samples with 4x and 80 samples with 8x. 6 groups indicates 50 samples with 35x, 30 samples with 50x, 20 samples with 100x, 90 samples with 4x, 80 samples with 8x and 30 samples with 3x. Results are based on 100,000 replicates for MAF=0.1. Base calling error is set to 0.01 across all groups. HRD: high read depth, LRD: low read depth.

Table S13. Empirical Type I error for joint rare variant analysis with respect to sample size with respect to sample size at the significant level of 0.05 between the vRVS method, a conventional score test using called genotypes (GC) and true genotype (true geno) with HRD and LRD groups for quantitative data analysis.

| Sample size | Type of analysis | | |
|---|---|---|---|
| | True geno (CAST/SKAT) | GC (CAST/SKAT) | vRVS (CAST/SKAT) |
| 100 | 0.0475/0.047 | 0.0505/0.0415 | 0.045/0.045 |
| 500 | 0.0515/0.047 | 0.0460/0.0490 | 0.0460/0.0470 |
| 1000 | 0.0475/0.0515 | 0.054/0.0535 | 0.0515/0.049 |

*Note:* 100 samples are composed of 4 groups; 20 samples with 35x, 40 samples with 50x, 25 samples with 4x and 15 samples with 8x. 500 samples are composed of 4 groups; 100 samples with 35x, 120 samples with 50x, 200 samples with 4x and 80 samples with 8x. 1000 samples are composed of 4 groups; 100 samples with 35x, 250 samples with 50x, 450 samples with 4x and 200 samples with 8x. Results are based on 10,000 replicates for MAF=0.01. vRVS is performed through bootstrap (1,000 iterations), GC and true genotype are performed through permutation (1,000 iterations). Each association test used a collapsed group of 5 variants. Base calling error is set to 0.01 across all groups. HRD: high read depth, LRD: low read depth.

Table S14. Empirical Type I error for joint rare variant analysis with respect to different number of groups combined at the significant level of 0.05 between the vRVS method, a conventional score test using called genotypes (GC) and true genotype (true geno) with HRD and LRD groups for quantitative data analysis.

| Number of groups combined | Type of analysis | | |
|---|---|---|---|
| | True geno (CAST/SKAT) | GC (CAST/SKAT) | vRVS (CAST/SKAT) |
| 2 groups | 0.0533/0.0522 | 0.0530/0.0520 | 0.5425/0.0483 |
| 3 groups | 0.0495/ 0.0507 | 0.0488/0.0542 | 0.0510/0.0515 |
| 4 groups | 0.0483/0.4583 | 0.0507/0.0530 | 0.0495/0.0482 |
| 5 groups | 0.0528/0.0458 | 0.0528/0.0523 | 0.0507/0.0455 |
| 6 groups | 0.0492/0.0520 | 0.0458/0.0502 | 0.0467/0.0443 |

*Note:* 2 groups indicates 100 samples with 35x and 200 samples with 4x. 3 groups indicates 50 samples with 35x, 50 samples with 50x and 200 samples with 4x. 4 groups indicates 50 samples with 35x, 50 samples with 50x, 120 samples with 4x and 80 samples with 8x. 5 groups indicates 50 samples with 35x, 30 samples with 50x, 20 samples with 100x, 120 samples with 4x and 80 samples with 8x. 6 groups indicates 50 samples with 35x, 30 samples with 50x, 20 samples with 100x, 90 samples with 4x, 80 samples with 8x and 30 samples with 3x. Results are based on 30,000 replicates for MAF=0.01. Base calling error is set to 0.01 across all groups. vRVS is performed through bootstrap (1,000 iterations), GC and true genotype are performed through permutation (1,000 iterations). Each association test used a collapsed group of 5 variants. HRD: high read depth, LRD: low read depth.

Table S15. Empirical Type I error for common variant analysis with respect to different correlation value between the genotype, G and the covariate Z at the significance level of 0.05 between the vRVS method, a conventional score test using called genotypes (GC) and true generating genotypes (true geno) with 100 HRD cases and 200 LRD control.

| Cor(G,Z) | Type of analysis | | |
|---|---|---|---|
| | True geno | GC | vRVS |
| 0 | 0.05045 | 0.05305 | 0.05087 |
| 0.01 | 0.05014 | 0.05214 | 0.04858 |
| 0.02 | 0.05049 | 0.05276 | 0.04894 |
| 0.05 | 0.05057 | 0.05259 | 0.04922 |
| 0.1 | 0.05072 | 0.05195 | 0.04789 |
| 0.2 | 0.05058 | 0.05572 | 0.04489 |
| 0.5 | 0.05184 | 0.05652 | 0.02471 |

*Note:* 100 cases are composed of 60 cases with 35x and 40 cases with 50x and 200 controls are composed of 50 controls with 4x and 150 controls with 8x. Results are based on 100,000 replicates for MAF=0.1. Base calling error  is set to 0.01 across all groups. HRD: high read depth, LRD: low read depth.

Table S16. Empirical Type I error for common variant analysis with respect to different correlation value between the genotype, G and the covariate Z at the significance level of 0.05 between the vRVS method, a conventional score test using called genotypes (GC) and true generating genotypes (true geno) with 500 HRD cases and 1500 LRD control.

| Cor(G,Z) | Type of analysis | | |
|---|---|---|---|
| | True geno | GC | vRVS |
| 0 | 0.0506 | 0.0630 | 0.0508 |
| 0.01 | 0.0504 | 0.0957 | 0.0502 |
| 0.02 | 0.0501 | 0.0946 | 0.0498 |
| 0.05 | 0.0505 | 0.0621 | 0.0507 |
| 0.1 | 0.0494 | 0.0780 | 0.0489 |
| 0.2 | 0.0509 | 0.0788 | 0.0473 |
| 0.5 | 0.0501 | 0.0622 | 0.0248 |

*Note:* 500 cases are composed of 200 cases with 35x and 300 cases with 50x and 1500 controls are composed of 600 controls with 4x and 900 controls with 8x. Results are based on 100,000 replicates for MAF=0.1. Base calling error  is set to 0.01 across all groups. HRD: high read depth, LRD: low read depth.

Table S17. Empirical Type I error for common variant analysis with respect to sample size at the significant level of 0.05 between the vRVS method, a conventional score test using called genotypes (GC) and true genotype (true geno) with HRD cases and LRD control.

| Sample size (case:control) | Type of analysis | | |
|---|---|---|---|
| | True geno | GC | vRVS |
| 100:50 | 0.0496 | 0.0516 | 0.0547 |
| 200:100 | 0.0502 | 0.0558 | 0.0511 |
| 1500:500 | 0.0505 | 0.0984 | 0.5205 |

*Note:* Cases are 35x and controls are with 4x. Results are based on 100,000 replicates for MAF=0.1. Base calling error is set to 0.01 across all groups. HRD: high read depth, LRD:low read depth, MAF=minor allele frequency.

Table S18. Empirical Type I error for common variant analysis with respect to sample sizes for cases and controls at the significant level of 0.05 between the vRVS method, a conventional score test using called genotypes (GC) and true genotype (true geno) with HRD cases and control.

| Sample size case:control | Type of analysis | | |
|---|---|---|---|
| | True geno | GC | vRVS |
| 50:50 | 0.0511 | 0.0511 | 0.0511 |
| 50:100 | 0.0512 | 0.0512 | 0.0512 |
| 100:50 | 0.0494 | 0.0494 | 0.0494 |
| 100:100 | 0.0490 | 0.0490 | 0.0490 |
| 100:200 | 0.0502 | 0.0502 | 0.0502 |
| 200:100 | 0.0506 | 0.0506 | 0.0506 |
| 250:50 | 0.0485 | 0.0485 | 0.0485 |
| 300:300 | 0.0499 | 0.0499 | 0.0499 |
| 300:600 | 0.0499 | 0.0499 | 0.0499 |
| 600:300 | 0.0506 | 0.0506 | 0.0506 |
| 500:1500 | 0.0498 | 0.0498 | 0.0498 |
| 1500:500 | 0.0495 | 0.0494 | 0.0494 |

*Note:* Cases are 32x and controls are with 50x. Results are based on 100,000 replicates for MAF=0.1. Base calling error is set to 0.01 across all groups.

Table S19. Empirical Type I error for common variant analysis with respect to sample sizes at the significant level of 0.05 between the vRVS method, a conventional score test using called genotypes (GC) and true genotype (true geno).

| Sample size Case:control | Type of analysis | | |
|---|---|---|---|
| | True geno | GC | vRVS |
| 50:50 | 0.0511 | 0.0536 | 0.0533 |
| 50:100 | 0.0500 | 0.0500 | 0.0552 |
| 100:50 | 0.0496 | 0.0557 | 0.0560 |
| 100:100 | 0.0508 | 0.0533 | 0.0517 |
| 100:200 | 0.0493 | 0.0510 | 0.0514 |
| 200:100 | 0.0498 | 0.0567 | 0.0533 |
| 250:50 | 0.0507 | 0.0577 | 0.0641 |
| 300:300 | 0.0501 | 0.0608 | 0.0508 |
| 300:600 | 0.0497 | 0.0617 | 0.0500 |
| 600:300 | 0.0497 | 0.0669 | 0.0511 |
| 500:1500 | 0.0491 | 0.0757 | 0.0492 |
| 1500:500 | 0.0498 | 0.0854 | 0.0498 |

*Note:* Cases are 4x and controls are with 5x. Results are based on 100,000 replicates for MAF=0.1. Base calling error is set to 0.01 across all groups.

Table S20. Empirical Type I error for common variant analysis with rwith unbalanced sample sizes for cases and controls at the significant level of 0.05 between the vRVS method, a conventional score test using called genotypes (GC) and true genotype (true geno).

| Sample size Case:control | Type of analysis | | |
|---|---|---|---|
| | True geno | GC | vRVS |
| 50:500 | 0.04913 | 0.06671 | 0.0474 |
| 200:1000 | 0.04985 | 0.08162 | 0.04865 |

*Note:* Cases are 35x and controls are with 4x. Results are based on 100,000 replicates for MAF=0.1. Base calling error is set to 0.01 across all groups.

Table S21. Empirical power for common variant analysis with unbalanced sample sizes for cases and controls at the significant level of 0.05 between the vRVS method,and true genotype (true geno) when the OR=1.5.

| Sample size Case:control | Type of analysis | |
|---|---|---|
| | True geno | vRVS |
| 50:500 | 0.28975 | 0.25799 |
| 200:1000 | 0.69461 | 0.6296 |

*Note:* Cases are 35x and controls are with 4x. Results are based on 100,000 replicates for MAF=0.1. Base calling error is set to 0.01 across all groups. OR:odds ratio

### 3 Derivations of the robust variance for the score test used in vRVS

#### 3.1 Quantitative trait analysis

##### 3.1.1 Calculation of the robust variance of the score for a common variant association test

- **No covariates**

  We derive the vRVS for the score test for quantitative trait association, where the quantitative trait is assumed to have a normal distribution. We also consider integrating sequence data from a arbitrary number of cohorts($k$). Our method can allow for covariate adjustment. Suppose we have two groups that we combine for a genetic association analysis where sequencing were done under different platforms. The score test for testing SNP $j$ is taking the same form under $H_0 : \beta_1 = 0$, which is $T_j = S_j^2/var(S_j)$ where $S_j = \sum_{i=1}^{n}(Y_i - E(Y_i))E(G_{ij} \mid D_{ij})$. For ease of notation, we drop $j$. Let $X_i = E(G_i \mid D_i)$.

  - Calculation of $var(S)$:

$$
\begin{aligned}
S &= \sum_{i=1}^{n}(Y_i - \bar{Y})X_i \\
Var(S) &= Var(\sum_{i=1}^{n}(Y_i - \bar{Y})X_i) \\
&= \sum_{i=1}^{n} Var((Y_i - \bar{Y})X_i) + \\
&\quad 2 \times \sum_{i=1}^{n-1} \sum_{j=i+1}^{n} Cov((Y_i - \bar{Y})X_i, (Y_j - \bar{Y})X_j)
\end{aligned}
$$

  Using $Var(XY) = Var(X)Var(Y) + E(X)^2Var(Y) + E(Y)^2Var(X)$ [Bohrnstedt and

Golderberger, 1969], we get

$$\sum_{i=1}^{n} Var((Y_i - \bar{Y})X_i) = \sum_{i=1}^{n}[Var(Y_i - \bar{Y})Var(X_i) +$$

$$E(X_i)^2 Var((Y_i - \bar{Y})) +$$

$$[E(Y_i - \bar{Y})]^2 Var(X_i)]$$

$$= (n-1)Var(Y_1)[Var(X_i) + (E(X_1))^2],$$

where $Var(Y_i - \bar{Y}) = Var(Y_i) + \frac{Var(Y_i)}{n}$ and $E(Y_i - \bar{Y}) = 0.$

The covariance part:

$$\begin{aligned}
Cov((Y_i - \bar{Y})X_i, (Y_j - \bar{Y})X_j) &= Cov(Y_i X_i, Y_j X_j) \\
&- Cov(\bar{Y} X_i, Y_j X_j) \\
&- Cov(Y_i X_i, \bar{Y} X_j) \\
&+ Cov(\bar{Y} X_i, \bar{Y} X_j) \\
&= 0 - \frac{1}{n} Cov(Y_j X_i, Y_j X_j)) \\
&- \frac{1}{n} Cov(Y_i X_i, Y_i X_j)) \\
&- \frac{1}{n^2} Cov(Y_k X_i, Y_k X_j)) \\
&= -\frac{1}{n} Cov(Y_k X_i, Y_k X_j))
\end{aligned}$$

$$2\sum_{i=1}^{n-1}\sum_{j=i+1}^{n} Cov((Y_i - \bar{Y})X_i, (Y_j - \bar{Y})X_j = -(n-1)E(X_1)^2 Var(Y_1)$$

Thus the variance of the score is:

$$
\begin{aligned}
Var(\sum_{i=1}^{n}(Y_i - \bar{Y})^2 X_i) &= (n-1)Var(Y)[Var(X_1) + (E(X_1))^2] \\
&\quad - (n-1)E(X_1)^2 Var(Y_1) \\
&= (n-1)Var(Y_1)Var(X_1)
\end{aligned}
$$

– Suppose $\underbrace{Y_i, Y_2, ..., Y_{n_1}}_{n_1}, \underbrace{Y_{n_1+1}, ..., Y_n}_{n_2} \sim N(\beta_0 + \beta_i g_i, \sigma^2), i = 1, ..., n$ where $n = n_1 + n_2$ and $n_1$ and $n_2$ are the sample sizes of HRD and LRD groups. Then the score and is its variance are as shown below:

$$
\begin{aligned}
S &= \sum_{i=1}^{n_1}(Y_i - \bar{Y})X_i + \sum_{i=n_1+1}^{n}(Y_i - \bar{Y})X_i \\
Var(S_j) &= \sum_{i=1}^{n_1} Var((Y_i - \bar{Y})X_i) \\
&\quad + \sum_{i=n_1+1}^{n} Var((Y_i - \bar{Y})X_i) \\
&\quad + 2 \times \sum_{i=1}^{n-1}\sum_{j=i+1}^{n} Cov((Y_i - \bar{Y})X_i, (Y_j - \bar{Y})X_j) \\
\sum_{i=1}^{n_1} Var((Y_i - \bar{Y})X_i) &= \sum_{i=1}^{n} Var(Y_i - \bar{Y})Var(X_i) + E(X_i)^2 Var(Y_i - \bar{Y}) \\
&= n_1 \frac{n-1}{n} Var(Y_1)[Var_1(X_{HRD}) + E(X_1)^2]
\end{aligned}
$$

$$Var(S) = n_1\frac{n-1}{n}Var(Y_1)Var_1(X_{HRD}) + n_2\frac{n-1}{n}Var(Y_1)Var_2(X_{LRD})$$
$$= \sum_{i=1}^{n}(Y_i - \bar{Y})^2[\frac{n_1}{n}Var_1(X_{HRD}) + \frac{n_2}{n}Var_2(X_{LRD})]$$

where $Var_1(X_{HRD})$ and $Var_2(X_{LRD})$ are estimated using the sample variance of $X$ in the HRD group and LRD group respectively. To add the robust adjustment, $Var_1(X_{HRD})$ will be replaced with $Var(G_{ij})$ which is estimated through the estimated $P(G_{ij} = g)$ using the full data set. With this estimated $P(G_{ij} = g)$ and under $H_0$, the expectation $E(X_{HRD}) = E(X_{LRD})$ [Derkach et al., 2014].

– If there are $K$ groups combined, the formula will look like:

$$Var(S) = \sum_{i=1}^{n}(Y_i - \bar{Y})^2[\sum_{k=1}^{K}\frac{n_k}{n}Var_k(X_k)]$$

where $Var_k(X_k)$ is the variance of $X$ for group $k$. It will be replaced with $Var(G_{ij})$ if it is a HRD group.

- **Covariates added**

Suppose $\underbrace{Y_i, Y_2, ..., Y_{n_1}}_{n_1}, \underbrace{Y_{n_1+1}, ..., Y_n}_{n_2} \sim N(\beta_0 + \beta_i g_i + \alpha z_i, \sigma^2)$, $i = 1, ..., n$ where $n = n_1 + n_2$ and $n_1$ and $n_2$ are the sample sizes of HRD and LRD groups. Then the score is: $S = \sum_{i=1}^{n}(Y_i - \hat{Y}_i)X_i + \sum_{i=n_1+1}^{n}(Y_i - \hat{Y}_i)X_i$, where $\hat{Y}_i$ are the fitted value under $H_0$. Note that $\hat{Y} = HY$, where H is the *hat* matrix, i.e. $H = Z(Z^TZ)^{-1}Z^T$, $e_i = (Y_i - \hat{Y}_i)$ and $Cov(\boldsymbol{e}) = (I - H)\sigma^2$, where $I_{n\times n}$ is the identity matrix with

$$Var(S) = \sum_{i=1}^{n_1}Var(e_iX_i) + \sum_{i=n_1+1}^{n}Var(e_iX_i) + \sum\sum_{i\neq j}Cov(e_iX_i, e_jX_j)$$

$$\sum_{i=1}^{n_1} Var(e_i X_i) = \sum_{i=1}^{n_1} Var(e_i)Var(X_i) + E[e_i]^2 Var(X_i) + E[X_i]^2 Var(e_i)$$
$$= \sum_{i=1}^{n_1} Var(e_i)[Var(X_i] + E[X_i]^2] \qquad \text{since } E(e_i) = 0$$
$$\sum\sum_{i \neq j} Cov(e_i X_i, e_j X_j) = \sum\sum_{i \neq j} E(e_i e_j)E(X_i)E(X_j)$$

Note that $E(e_i e_j) = Cov(e_i e_j) = \sigma^2[I - H][i, j]$ and $[I - H][i, j]$ is the $(i, j)^{th}$ entry of the matrix $(I - H)$. So $\sum\sum_{i \neq j}[I - H][i, j]$ is the sum of all off diagonal elements.

$$\sum\sum_{i \neq j} Cov(e_i X_i, e_j X_j) = -(n - k)\sigma^2 E[X_1]^2 \qquad (E(X_i) = E(X_j))$$

where $k$ is the number of columns in $H$ (number of covariates plus 1). Note that $Var(e_i) = \sigma^2(I - H)[i, i]$ and $\sum_{i=1}^{n_1} Var(e_i) = \sum_{i=1}^{n_1} \sigma^2(I - H)[i, i] = \sigma^2(n_1 - k_1)$, where $k_1$ is the sum of diagonals of the matrix $H$ for the HRD group, i.e. sum of the diagonals in $H_{n_1 \times n_1}$.

$$Var(S) = \sigma^2(n_1 - k_1)Var_1(X_{HRD}) + \sigma^2(n_1 - k_1)E[X_1]^2 + \sigma^2(n_2 - k_2)Var_2(X_{LRD})$$
$$+ \sigma^2(n_2 - k_2)E[X_1]^2 - \sigma^2(n - k)E[X_1]^2$$
$$= \sigma^2[(n_1 - k_1)Var_1(X_{HRD}) + (n_2 - k_2)Var_2(X_{LRD})]$$
$$= \sum_{i=1}^{n}(Y_i - \hat{Y}_i)^2[\frac{n_1 - k_1}{n - k}Var_1(X_{HRD}) + \frac{n_2 - k_2}{n - k}Var_2(X_{LRD})]$$
$$\approx \sum_{i=1}^{n}(Y_i - \hat{Y}_i)^2[\frac{n_1}{n}Var_1(X_{HRD}) + \frac{n_2}{n}Var_2(X_{LRD})]$$

Note that $\hat{\sigma^2} = \sum_{i=1}^{n}(Y_i - \hat{Y}_i)^2/(n - k)$. $Var_1(X_{HRD})$ and $Var_2(X_{LRD})$ are estimated using the sample variance of $X$ in the HRD group and LRD group respectively.

- If there are $K$ groups combined, the formula will look like:

$$Var(S) = \sum_{i=1}^{n}(Y_i - \hat{Y}_i)^2[\sum_{k=1}^{K}\frac{n_k}{n}Var_k(X_k)]$$

where $Var_k(X_k)$ is the variance of $X_i$ for group $k$. It will be replaced with $Var(G_{ij})$ if it is a HRD group.

##### 3.1.2 Calculation of the robust variance of the score for a joint rare variant association test

Suppose two variants are collapsed. Then

$$\boldsymbol{S} = [\underbrace{\sum_{i=1}^{n_1}(Y_i - \hat{Y}_i)X_i + \sum_{i=n_1+1}^{n}(Y_i - \hat{Y}_i)X_i}_{A} \quad \underbrace{\sum_{j=1}^{n_1}(Y_j - \hat{Y}_j)Z_j + \sum_{j=n_1+1}^{n}(Y_j - \hat{Y}_j)Z_j}_{B}]$$

$$\boldsymbol{Cov(S)} = \begin{bmatrix} Var(A) & Cov(A,B) \\ Cov(A,B) & Var(B) \end{bmatrix}$$

$Var(A)$ was shown in Section 3.2.1.

$$
\begin{aligned}
Cov(A,B) = Cov(\sum_{i=1}^{n_1} e_i X_i, \sum_{j=1}^{n_1} e_i Z_j) \qquad & ① \\
+ Cov(\sum_{i=1}^{n_1} e_i X_i, \sum_{j=n_1+1}^{n} e_i Z_j) \qquad & ② \\
+ Cov(\sum_{i=n_1+1}^{n} e_i X_i, \sum_{j=1}^{n_1} e_i Z_j) \qquad & ③ \\
+ Cov(\sum_{i=n_1+1}^{n} e_i X_i, \sum_{j=n_1+1}^{n} e_i Z_j) \qquad & ④
\end{aligned}
$$

$$
\begin{aligned}
① \quad Cov(\sum_{i=1}^{n_1} e_i X_i, \sum_{j=1}^{n_1} e_j Z_j) &= \underbrace{\sum_{i=1}^{n_1}[E(e_i^2)E(X_i Z_i) - E[e_i]^2 E(X_i)E(Z_i)]}_{i=j} \\
&+ \underbrace{\sum_{i=1}^{n_1}\sum_{j=1}^{n_1} E(X_i)E(Z_i)Cov(e_i, e_j)}_{i \neq j}
\end{aligned}
$$

$$
② \quad Cov(\sum_{i=1}^{n_1} e_i X_i, \sum_{j=n_1+1}^{n} e_j Z_j) = \sum_{i=1}^{n_1}\sum_{j=n_1+1}^{n} E(X_i)E(Z_j)Cov(e_i, e_j)
$$

$$
③ \quad Cov(\sum_{i=n_1+1}^{n} e_i X_i, \sum_{j=1}^{n_1} e_j Z_j) = \sum_{i=n_1+1}^{n}\sum_{j=1}^{n_1} E(X_i)E(Z_j)Cov(e_i, e_j)
$$

$$④ \quad Cov(\sum_{i=n_1+1}^{n} e_i X_i, \sum_{j=n_1+1}^{n} e_j Z_j) = \underbrace{\sum_{i=n_1+1}^{n} [E(e_i^2)E(X_i Z_i) - E[e_i]^2 E(X_i)E(Z_i)]}_{i=j}$$

$$+ \underbrace{\sum_{i=n_1+1}^{n} \sum_{j=n_1+1}^{n} E(X_i)E(Z_i)Cov(e_i, e_j)}_{i \neq j}$$

Note that,

$$① = \sum_{i=1}^{n_1} [E(e_i^2)[E(X_i Z_i) - E(X_i)E(Z_i)] + E(e_i^2)E(X_i)E(_i) - E[e_i]^2 E(X_i)E(Z_i)]$$

$$+ \underbrace{\sum_{i=1}^{n_1} \sum_{j=1}^{n_1}}_{i \neq j} E(X_i)E(Z_i)Cov(e_i, e_j)$$

$$= \sum_{i=1}^{n_1} [E[e_i^2] Cov_1(X_{HRD}, Z_{HRD}) + Var(e_i)E(X_i)E(Z_i)] + \underbrace{\sum_{i=1}^{n_1} \sum_{j=1}^{n_1}}_{i \neq j} E(X_i)E(Z_i)Cov(e_i, e_j)$$

$$④ = \sum_{i=n_1+1}^{n} [E[e_i^2] Cov_2(LRD, LRD) + Var(e_i)E(X_i)E(Z_i)] + \underbrace{\sum_{i=n_1+1}^{n} \sum_{j=n_1+1}^{n}}_{i \neq j} E(X_i)E(Z_i)Cov(e_i, e_j)$$

$$② + ③ = \sum_{i=1}^{n_1} \sum_{j=n_1+1}^{n} E(X_i)E(Z_j)Cov(e_i, e_j) + \sum_{i=n_1+1}^{n} \sum_{j=1}^{n_1} E(X_i)E(Z_j)Cov(e_i, e_j)$$

$$
\begin{aligned}
① + ② + ③ + ④ = &\sum_{i=1}^{n_1} E[e_i^2]Cov_1(X_{HRD}, Z_{HRD}) + \sum_{i=n_1+1}^{n} E[e_i^2]Cov_2(X_{LRD}, Z_{LRD}) \\
&+ \sum_{i=1} E(X_i)E(Z_i)Var(e_i) + \sum_{i=n_1+1}^{n} E(X_i)E(Z_i)Var(e_i) \\
&+ \underbrace{\sum_{i=1}^{n_1}\sum_{j=1}^{n_1}}_{i \neq j} E(X_i)E(Z_i)Cov(e_i, e_j) + \underbrace{\sum_{i=n_1+1}^{n}\sum_{j=n_1+1}^{n}}_{i \neq j} E(X_i)E(Z_i)Cov(e_i, e_j) \\
&+ ② + ③
\end{aligned}
$$

The last six terms in the equation above equals to $E(X_1)E(Z_1)\sigma^2 \sum_{i=1}^{n}\sum_{j=1}^{n}(I - H)[i,j] = 0$ since sum of all elements of $(I - H)$ is 0 Then

$$
\begin{aligned}
Cov(\boldsymbol{S}) &= \sigma^2 \sum_{i=1}^{n_1}(1 - H)[i,i]Cov_1(X_{HRD}, Z_{HRD}) + \sum_{i=n_1+1}^{n}(1 - H)[i,i]Cov_2(X_{LRD}, Z_{LRD}) \\
&= \sum_{i=1}^{n}(Y_i - \hat{Y}_i)^2[\frac{n_1 - k_1}{n - k}Cov_1(X_{HRD}, Z_{HRD}) + \frac{n_2 - k_2}{n - k}Cov_2(X_{LRD}, Z_{LRD})] \\
&\approx \sum_{i=1}^{n}(Y_i - \hat{Y}_i)^2[\frac{n_1}{n}Cov_1(X_{HRD}, Z_{HRD}) + \frac{n_2}{n}Cov_2(X_{LRD}, Z_{LRD})]
\end{aligned}
$$

$Cov_1(X_{HRD}, Z_{HRD})$ is replaced with the robust variance estimate. $Cov_1(X_{HRD}, Z_{HRD}) = V^T cor(X_{HRD}, Z_{HRD})V$, where $V$ is a diagonal matrix with diagonals $\sqrt{Var(G_{ij})}$ for the corresponding collapsed variants.

#### 3.2 Binary trait analysis

The variance of the score, $S = \sum_{i=1}^{n}(Y - \bar{Y})E(G_i \mid D_i)$ when there are no covariates was shown in Methods section and in [Derkach et al., 2014]. Again for ease of notation let $X_i = E(G_i \mid D_i)$ and $n_1$ the number of cases (sequenced at high read depth), $n_2$ the number of controls (sequenced at low read depth) and $n = n_1 + n_2$.

##### 3.2.1 Calculation of the robust variance of the score for a common variant association test

- **Covariates added**

We calculate the variance as following

$$
\begin{aligned}
S &= \sum_{i=1}^{n_1} (Y_i - \hat{Y}_i)X_i + \sum_{i=n_1+1}^{n} (Y_i - \hat{Y}_i)X_i \\
Var(S) &= \sum_{i=1}^{n_1} Var((1 - \hat{Y}_i)X_i) + \sum_{i=n_1+1}^{n} Var((-\hat{Y}_i)X_i) \\
&+ 2 \times \sum_{i=1}^{n_1-1} \sum_{j=i+1}^{n_1} Cov((1 - \hat{Y}_i)X_i, (1 - \hat{Y}_j)X_j) \\
&+ 2 \times \sum_{i=1}^{n_2-1} \sum_{j=i+1}^{n_2} Cov((-\hat{Y}_i)X_i, (-\hat{Y}_j)X_j) \\
&+ 2 \times \sum_{i=1}^{n_1} \sum_{j=i+1}^{n_2} Cov((1 - \hat{Y}_i)X_{ij}, (-\hat{Y}_i)X_j)
\end{aligned}
$$

Since $Cov((1-\hat{Y}_i)X_i, (1-\hat{Y}_j)X_j) = Cov((-\hat{Y}_i)X_i, (-\hat{Y}_j)X_j) = Cov((1-\hat{Y}_i)X_{ij}, (-\hat{Y}_i)X_j) = Cov(\hat{Y}_i)X_i, \hat{Y}_j)X_j) = E(X_i)E(X_j)Cov(\hat{Y}_i, \hat{Y}_j)$, summation of all covariance terms would be $2\sum_{i=1}^{n-1}\sum_{j=i+1}^{n} E(X_i)E(X_j)Cov(\hat{Y}_i, \hat{Y}_j)$

$$\sum_{i=1}^{n_1} Var((1-\hat{Y}_i)X_i) = \sum_{i=1}^{n_1}\{Var(1-\hat{Y}_i)Var(X_i) + E(X_i)^2 Var(1-\hat{Y}_i) + (E(1-\hat{Y}_i))^2 Var(X_i)\}$$

$$\sum_{i=1}^{n_1} Var((\hat{Y}_i)X_i) = \sum_{i=1}^{n_2}\{Var(-\hat{Y}_i)Var(X_i) + E(X_i)^2 Var(-\hat{Y}_i) + (E(-\hat{Y}_i))^2 Var(X_i)\}$$

Note that:

$$\sum_{i=1}^{n_1} E[(1-\hat{Y}_i)^2]Var(X_i) = \sum_{i=1}^{n_1}\{Var(1-\hat{Y}_i)Var(X_i) + (E(1-\hat{Y}_i))^2 Var(X_i)\}$$

$$\sum_{i=1}^{n_2} E[(-\hat{Y}_i)^2]Var(X_i) = \sum_{i=1}^{n_2}\{Var(-\hat{Y}_i)Var(X_i) + (E(-\hat{Y}_i))^2 Var(X_i)\}$$

$$\sum_{i=1}^{n} E(X_i)^2 Var(\hat{Y}_i) = \sum_{i=1}^{n_1} E(X_i)^2 Var(1-\hat{Y}_i)) + \sum_{i=1}^{n_1} E(X_i)^2 Var(-\hat{Y}_i))$$

Thus,

$$Var(S) = \sum_{i=1}^{n_1} E[(1-\hat{Y}_i)^2]Var(X_i) + \sum_{i=1}^{n_2} E[(-\hat{Y}_i)^2]Var(X_i) + \sum_{i=1}^{n} E(X_i)^2 Var(\hat{Y}_i) + 2\sum_{i=1}^{n-1}\sum_{j=i+1}^{n} E(X_i)E(X_j)Cov(\hat{Y}_i, \hat{Y}_j)$$

Under $H_0$, the expectation $E(X_i) = E(X_j)$ because the mean of $E(G_{ij}|D_{ij})$ is equal for cases and controls, thus

$$\sum_{i=1}^{n} E(X_i)^2 Var(\hat{Y}_i) + 2\sum_{i=1}^{n-1}\sum_{j=i+1}^{n} E(X_i)E(X_j)Cov(\hat{Y}_i,\hat{Y}_j) = E(X_1)^2 Var(\hat{Y}_1 + \hat{Y}_2 + ... + \hat{Y}_n)$$
$$= 0$$

since $\sum_{i=1}^{n} \hat{Y}_i = \sum_{i=1}^{n} Y_i = n_1$(number of cases).

We estimate $E[(1-\hat{Y}_i)^2]$ by $\sum_{i_1}^{n_1}(1-\hat{Y}_i)^2/n_1$ and $E[(-\hat{Y}_i)^2]$ by $\sum_{n_1+1}^{n}(-\hat{Y}_i)^2/n_2$. Then

$$\hat{V}ar(S) = \sum_{i=1}^{n_1}(1-\hat{Y}_i)^2\hat{V}ar_{case}(X_{case})) + \sum_{i=n_1+1}^{n_2}(-\hat{Y}_i)^2\hat{V}ar_{cont}(X_{cont}))$$

where $\hat{V}ar_{case}(X_{case})$ is the sample variance of X in case group and $\hat{V}ar_{cont}(X_{cont})$ is the sample variance of X in case group. $Var_{case}(X_{case})$ is replaced with $Var(G_{ij})$ which is estimated through the estimated $P(G_{ij} = g)$ using the full data set.

##### 3.2.2 Calculation of the robust variance of the score for a joint rare variant association test

- **Covariates added**

Suppose two variants are collapsed. Then

$$\boldsymbol{S} = [\underbrace{\sum_{i=1}^{n_1}(Y_i-\hat{Y}_i)X_i + \sum_{i=n_1+1}^{n}(Y_i-\hat{Y}_i)X_i}_{A} \quad \underbrace{\sum_{j=1}^{n_1}(Y_j-\hat{Y}_j)Z_j + \sum_{j=n_1+1}^{n}(Y_j-\hat{Y}_j)Z_j}_{B}]$$

$$\boldsymbol{Cov(S)} = \begin{bmatrix} Var(A) & Cov(A,B) \\ Cov(A,B) & Var(B) \end{bmatrix}$$

$Var(A)$ was shown in Section 3.2.1.

$$
\begin{aligned}
Cov(A,B) = {} & Cov(\sum_{i=1}^{n_1}(1-\hat{Y}_i)X_i, \sum_{j=1}^{n_1}(1-\hat{Y}_j)Z_j) & ① \\
& + Cov(\sum_{i=1}^{n_1}(1-\hat{Y}_i)X_i, \sum_{j=n_1+1}^{n}(-\hat{Y}_j)Z_j) & ② \\
& + Cov(\sum_{i=n_1+1}^{n}(-\hat{Y}_i)X_i, \sum_{j=1}^{n_1}(1-\hat{Y}_j)Z_j) & ③ \\
& + Cov(\sum_{i=n_1+1}^{n}(-\hat{Y}_i)X_i, \sum_{j=n_1+1}^{n}(-\hat{Y}_j)Z_j) & ④
\end{aligned}
$$

$$
\begin{aligned}
① \quad Cov(\sum_{i=1}^{n_1}(1-\hat{Y}_i)X_i, \sum_{j=1}^{n_1}(1-\hat{Y}_j)Z_j) = {} & \sum_{i=1}^{n_1}\sum_{j=1}^{n_1} Cov(X_i, Z_i) - Cov(X_i, \hat{Y}_j Z_j) \\
& - Cov(\hat{Y}_i X_i, Z_i) + Cov(\hat{Y}_i X_i, \hat{Y}_j Z_i)
\end{aligned}
$$

$$
\begin{aligned}
① = {} & \underbrace{\sum_{i=1}^{n_1} Cov(X_i, X_i) - 2E(\hat{Y}_i)Cov(X_i, X_i) + [E(\hat{Y}_i^{\,2})E(X_i Z_i) - E(\hat{Y}_i)^2 E(X_i)E(Z_i)]}_{i=j} \\
& + \underbrace{\sum_{i=1}^{n_1}\sum_{j=1}^{n_1} E(X_i)E(Z_j)Cov(\hat{Y}_i, \hat{Y}_j)}_{i \neq j}
\end{aligned}
$$

$$② \quad Cov(\sum_{i=1}^{n_1}(1-\hat{Y}_i)X_i, \sum_{j=n_1+1}^{n}(-\hat{Y}_j)Z_j) = \sum_{i=1}^{n_1}\sum_{j=n_1+1}^{n} E(X_i)E(Z_j)Cov(\hat{Y}_i, \hat{Y}_j)$$

$$③ \quad Cov(\sum_{i=n_1+1}^{n}(-\hat{Y}_i)X_i, \sum_{j=1}^{n_1}(1-\hat{Y}_j)Z_j) = \sum_{i=n_1+1}^{n}\sum_{j=1}^{n_1} E(X_i)E(Z_j)Cov(\hat{Y}_i, \hat{Y}_j)$$

$$④ \quad Cov(\sum_{i=n_1+1}^{n}(-\hat{Y}_i)X_i, \sum_{j=n_1+1}^{n}(-\hat{Y}_j)Z_j) = \sum_{i=n_1+1}^{n}\sum_{j=n_1+1}^{n} E(X_i)E(Z_j)Cov(\hat{Y}_i, \hat{Y}_j)$$

$$④ = \underbrace{\sum_{i=n_1+1}^{n} E(\hat{Y}_i^{\,2})E(X_iZ_i) - E(\hat{Y}_i)^2E(X_i)E(Z_i)}_{i=j}$$
$$+ \underbrace{\sum_{i=1}^{n_1}\sum_{j=1}^{n_1} E(X_i)E(Z_j)Cov(\hat{Y}_i, \hat{Y}_j)}_{i\neq j}$$

Before we add ①+②+③+④, note that,

$$
\begin{aligned}
① = &\sum_{i=1}^{n_1} E[(1-\hat{Y}_i)^2]Cov_{case}(X,Z) - \sum_{i=1}^{n_1} E[(\hat{Y}_i)^2]Cov_{case}(X,Z) + \sum_{i=1}^{n_1} E[(\hat{Y}_i)^2]E_{case}(XZ) \\
&- \sum_{i=1}^{n_1} E[(\hat{Y}_i)]^2 E(X_i)E(Z_i) + \underbrace{\sum_{i=1}^{n_1}\sum_{j=1}^{n_1}}_{i\neq j} E(X_i)E(Z_i)Cov(\hat{Y}_i,\hat{Y}_j) \\
= &\sum_{i=1}^{n_1} E[(1-\hat{Y}_i)^2]Cov_{case}(X,Z) + \sum_{i=1}^{n_1} E[(\hat{Y}_i)^2]E(X_i)E(Z_i) \\
&- \sum_{i=1}^{n_1} E([\hat{Y}_i)]^2 E(X_i)E(Z_i) + \underbrace{\sum_{i=1}^{n_1}\sum_{j=1}^{n_1}}_{i\neq j} E(X_i)E(Z_i)Cov(\hat{Y}_i,\hat{Y}_j) \\
= &\sum_{i=1}^{n_1} E[(1-\hat{Y}_i)^2]Cov_{case}(X,Z) + \sum_{i=1}^{n_1} Var(\hat{Y}_i)E(X_i)E(Z_i) + \underbrace{\sum_{i=1}^{n_1}\sum_{j=1}^{n_1}}_{i\neq j} E(X_i)E(Z_i)Cov(\hat{Y}_i,\hat{Y}_j)
\end{aligned}
$$

$$
\begin{aligned}
④ = &\sum_{i=n_1+1}^{n} E[(\hat{Y}_i)^2]Cov_{cont}(X,Z) + \sum_{i=n_1+1}^{n} E[(\hat{Y}_i)^2]E(X_i)E(Z_i) - \sum_{i=n_1+1}^{n} E[(\hat{Y}_i)]^2 E(X_i)E(Z_i) \\
&+ \underbrace{\sum_{i=n_1+1}^{n}\sum_{j=n_1+1}^{n}}_{i\neq j} E(X_i)E(Z_i)Cov(\hat{Y}_i,\hat{Y}_j) \\
= &\sum_{i=n_1+1}^{n} E[(\hat{Y}_i)^2]Cov_{cont}(X,Z) + \sum_{i=n_1+1}^{n} Var(\hat{Y}_i)E(X_i)E(Z_i) \\
&+ \underbrace{\sum_{i=n_1+1}^{n}\sum_{j=n_1+1}^{n}}_{i\neq j} E(X_i)E(Z_i)Cov(\hat{Y}_i,\hat{Y}_j)
\end{aligned}
$$

Then

$$
\begin{aligned}
① + ② + ③ + ④ &= \sum_{i=1}^{n_1} E[(1-\hat{Y}_i)^2] Cov_{case}(X,Z) + \sum_{i=n_1+1}^{n} E[(\hat{Y}_i)^2] Cov_{cont}(X,Z) \\
&\quad + E(X_1)E(Z_1)Var(\hat{Y}_1 + \hat{Y}_2 + ... + \hat{Y}_n) \\
&= \sum_{i=1}^{n_1} E[(1-\hat{Y}_i)^2] Cov_{case}(X,Z) + \sum_{i=n_1+1}^{n} E[(\hat{Y}_i)^2] Cov_{cont}(X,Z)
\end{aligned}
$$

since $Var(\hat{Y}_1 + \hat{Y}_2 + ... + \hat{Y}_n) = Var(n_{case}) = 0$.

We estimate $E[(1-\hat{Y}_i)^2$ by $\sum_{i=1}^{n_1}(1-\hat{Y}_i)^2/n_1$ and $E[(-\hat{Y}_i)^2]$ by $\sum_{i=n_1+1}^{n}(-\hat{Y}_i)^2/n_2$. Then

$$
\hat{C}ov(\boldsymbol{S}) = \sum_{i=1}^{n_1}(1-\hat{Y}_i)^2 \hat{C}ov_{case}(X_{case}, Z_{case}) + \sum_{i=n_1+1}^{n} (-\hat{Y}_i)^2 \hat{C}ov_{cont}(X_{cont}, Z_{cont}))
$$

Again $Cov_{case}(X_{case}, Z_{case}) = V^T cor(X_{case}, Z_{case}) V$, where $V$ is a diagonal matrix with diagonals $\sqrt{Var(G_{ij})}$ for the corresponding collapsed variants.
